# Comprehensive analyses of partially methylated domains and differentially methylated regions in esophageal cancer reveal both cell-type- and cancer-specific epigenetic regulation

**DOI:** 10.1101/2022.08.18.504390

**Authors:** Yueyuan Zheng, Benjamin Ziman, Allen S. Ho, Uttam K. Sinha, Li-Yan Xu, En-Min Li, H Phillip Koeffler, Benjamin P. Berman, De-Chen Lin

**Affiliations:** Clinical Big Data Research Center, Scientific Research Center, The Seventh Affiliated Hospital of Sun Yat-sen University, Shenzhen 518107, P.R. China; Department of Medicine, Samuel Ochin Comprehensive Cancer Institute, Cedars-Sinai Medical Center, Los Angeles, USA; Center for Craniofacial Molecular Biology, Herman Ostrow School of Dentistry, and Norris Comprehensive Cancer Center, University of Southern California, Los Angeles, CA 90033, USA; Division of Otolaryngology-Head and Neck Surgery, Department of Surgery, Samuel Oschin Cancer Center, Cedars-Sinai Medical Center, Los Angeles, CA, USA; Department of otolaryngology, Keck School of Medicine, University of Southern California; The Key Laboratory of Molecular Biology for High Cancer Incidence Coastal Chaoshan Area, Shantou University Medical College, Guangdong, China; Department of Developmental Biology and Cancer Research, Institute for Medical Research Israel-Canada, Faculty of Medicine, The Hebrew University of Jerusalem, Israel

## Abstract

As one of the most common malignancies, esophageal cancer has two subtypes, squamous cell carcinoma (ESCC) and adenocarcinoma (EAC), arising from distinct cells-of-origin. However, distinguishing cell-type-specific molecular features from cancer-specific characteristics has been challenging. Here, we analyze whole-genome bisulfite sequencing (WGBS) data on 45 esophageal tumor and nonmalignant samples from both subtypes. We develop a novel sequence-aware method to identify large partially methylated domains (PMDs), revealing profound heterogeneity at both the methylation level (depth) and genomic distribution (breadth) of PMDs across tumor samples. We identify subtype-specific PMDs, which are associated with repressive transcription, chromatin B compartments and high somatic mutation rate. While the genomic locations of these PMDs are pre-established in normal cells, the degree of loss is significantly higher in tumors. We find that cell-type-specific deposition of H3K36me2 may underlie the genomic distribution PMDs. At a smaller genomic scale, both cell-type- and cancer-specific differentially methylated regions (DMRs) are identified for each subtype. Using binding motif analysis within these DMRs, we show that a cell-type-specific transcription factor such as HNF4A can maintain the binding sites that it establishes in normal cells, while being recruited to new binding sites with novel partners such as FOSL1 in cancer. Finally, leveraging pan-tissue single-cell and pan-cancer epigenomic datasets, we demonstrate that a substantial fraction of the cell-type-specific PMDs and DMRs identified here in esophageal cancer, are actually markers that co-occur in other cancers originating from related cell types. These findings advance our understanding of the DNA methylation dynamics at various genomic scales in normal and malignant states, providing novel mechanistic insights into cell-type- and cancer-specific epigenetic regulations.

## Introduction

Ranking seventh in cancer incidence and sixth in mortality worldwide, esophageal carcinoma is highly aggressive and its patients have poor outcomes, with a 5-year survival rate lower than 20%^1,2^. Esophageal cancer comprises two major histologic subtypes: squamous cell carcinoma (ESCC) and adenocarcinoma (EAC). These two subtypes have distinct clinical characteristics. ESCC occurs predominantly in the upper and mid-esophagus; EAC is prevalent in the lower esophagus near the gastroesophageal junction (GEJ) and is associated with the precursor lesion known as Barrett’s esophagus (BE). Biologically, ESCC arises from the squamous epithelial cells and has common features with other squamous cell carcinomas (SCC), such as head and neck SCC (HNSCC). In comparison, EAC has columnar cell features and shares many characteristics with tubular gastrointestinal adenocarcinomas. In particular, EAC is almost indistinguishable from GEJ adenocarcinoma in terms of genomic, biological and clinical features.

Epigenetically, multiple studies have reported molecular changes in esophageal cancer, especially at the DNA methylation level^3–9^. For example, methylation differences across thousands of loci between ESCC and EAC were noted by The Cancer Genome Atlas (TCGA)^3^ consortium. However, these prior works focused largely on the analyses of DNA methylation in gene promoter regions, which only make up ~6% of all CpG sites across the human genome. DNA methylation is known to play important roles in other noncoding regions, such as enhancers^10^, partially methylated domains (PMDs)^11^, as well as repetitive elements^12^. Therefore, the DNA methylome of esophageal cancer awaits further and comprehensive characterization through genome-wide single-base resolution approaches such as whole-genome bisulfite sequencing (WGBS).

CpG island (CGI) promoter hypermethylation and global DNA hypomethylation are two epigenomic hallmarks in cancer^13^. In most healthy tissues, the vast majority of CpG sites (>80%) across the genome are fully methylated, except for the CpG-rich regions (e.g., CGIs) and other regulatory elements (predominantly enhancers)^14^. Indeed, focal demethylation is a reliable signature of gene promoters and enhancers, and their methylation levels are robustly maintained across healthy tissues. Additionally, methylation patterns of CpG sites across the genome are notably variable across various normal cell types, and can be grouped into cell-type-specific differentially methylated regions (DMRs), which are linked to cell-type-specific regulatory regions^14,15^. By contrast, abnormal CGI promoter hypermethylation is frequently observed in cancer, which is commonly associated with long-term and stable gene repression^14^.

With respect to the global methylation loss, large hypomethylated blocks, also known as PMDs, cover more than one-third of the genome and coincide with heterochromatin, chromatin “B” compartment (determined by HiC) and nuclear lamina associated domains^16–18^. We and others recently found that accumulation of PMD hypomethylation is linked to cumulative mitotic cell divisions, late replication timing as well as the deposition of the histone mark H3K36me3^19,20^. Functionally, PMDs are associated with inactive gene transcription, heightened genomic instability and may be accompanied by activation of transposable elements (TEs)^19,21^. While incompletely understood, the majority of the PMD regions are possibly shared across developmental lineages^19^. However, there are enough cell-type specific PMDs to differentiate between different cancer cell types^17,22,23^ and between different healthy cell types^24^.

Several important questions on cell-type- and cancer-specific DMRs and PMDs await further characterization, including: i) the degree of the regional specificity of these domains (i.e, the proportions of DMR/PMD that are cell-type- and cancer-specific), ii) the functional significance of DMRs and PMDs in cancer biology, and iii) underlying mechanisms of the alteration of DMRs and PMDs during tumorigenesis. To address these questions, we performed analyses of WGBS data generated from a cohort of 45 esophageal samples, including 21 ESCC and 5 nonmalignant esophageal squamous (NESQ) tissues, as well as 12 EAC/GEJ tumors and 7 nonmalignant GEJ (NGEJ) tissues (**Fig. 1A**). We chose esophageal cancer as the disease model considering that the two subtypes are developed from distinct cell-of-origins, and we hypothesized that characterization of their methylome profiles might reveal cell-type- and cancer-specific methylation changes, together with underlying epigenetic mechanisms.

**Figure 1.**
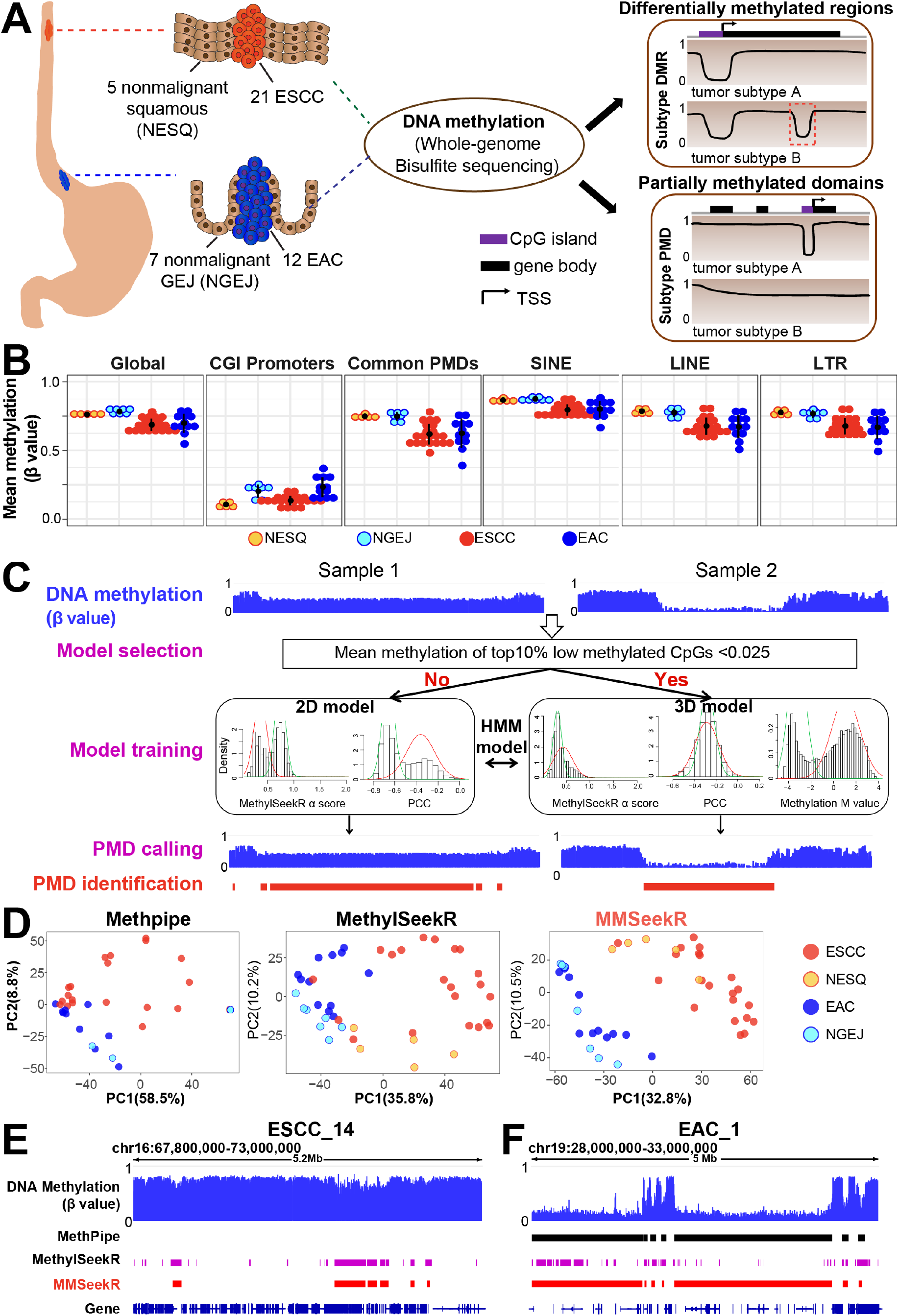
Identification of PMDs in esophageal samples by a sequence-aware multi-model PMD caller (MMSeekR). **(A)** A graphic model of the present study design. **(B)** Dot plots showing average methylation levels for all CpGs across the whole genome, CpGs within CGI promoters, common PMDs, SINE, LINE and LTR in different samples. The annotations from Takai et al^62^. were used for CGI methylation quantification. **(C)** Development of a new PMD caller. The MethylSeekR α score measures the distribution of methylation levels in sliding windows with 201 consecutive CpGs across the genome. α score < 1 corresponds to a polarized distribution towards a high or low methylation level (that is, HMDs), while α score >=1 corresponds to the distribution towards intermediate methylation levels (that is, PMDs). PCC shows the correlation between the predicted hypomethylation score based on a NN model, and the actual methylation level. A strong negative correlation indicates regions favoring PMDs, while weak/null correlation favors HMDs. **(D)** PCA analysis of 45 esophageal samples using the top 5,000 most variable 30-kb tiles for the three PMD callers. **(E-F)** Representative windows showing PMDs successfully identified by MMSeekR but failed to be detected by either MethPipe **(E)** or MethylSeekR **(F)**.

## Results

### Development of a novel sequence-aware calling method to identify PMDs

To characterize the esophageal cancer methylome, we analyzed WGBS profiles of 45 esophageal samples from two different cancer subtypes and their corresponding nonmalignant tissues^25^ (**Fig. 1A, Supplementary Fig. 2A**). All of the nonmalignant esophageal squamous (NESQ) tissues showed high inter-sample correlation despite that they were from two different cohorts (**Supplementary Fig. 2B** and **Supplementary Table 1**). To analyze the overall methylation pattern, we first investigated the methylation level at various genomic domains (**Fig. 1B**). As anticipated, both global hypomethylation (especially in common PMDs, defined as shared PMDs identified from 40 different cancer types^19^) and CGI promoter hypermethylation were observed in tumor samples. EAC tumors harbored notably higher methylation levels in CGI promoters than ESCC tumors, in line with TCGA results showing that gastrointestinal adenocarcinoma had higher frequency of CGI hypermethylation than cancers from most other tissues^26^. Interestingly, most NGEJ tissues showed higher CGI promoter methylation levels than NESQ tissues, and usually even higher than ESCC tumor samples. Similar to EAC, BE samples (a recognized precursor lesion of EAC) were reported to have a hypermethylation pattern at CGI promoters^7^. Since our NGEJ tissues were pathologically confirmed as inflammatory tissues but devoid of apparent BE, this result suggests that CGI hypermethylation may occur in inflamed GEJ. Interestingly, CGI hypermethylation has been observed in long-term-cultured colon organoids and cells upon prolonged exposure to cigarette smoke extract^27,28^. These data suggest that prolonged extrinsic pressure may result in DNA methylation changes at CGIs. Repetitive elements, especially from the LINE and LTR classes, lost DNA methylation in tumors compared with nonmalignant tissues (**Fig. 1B**), which might be accompanied with the activation of repetitive elements in tumor samples^21,29^.

Considering the importance of PMDs in cancer biology^17,19,22,23^, we sought to characterize this epigenomic domain in depth. Computational tools have been developed for the identification of PMDs, including MethPipe^30^ and MethylSeekR^31^. However, they sometimes fail or return unsatisfactory results for WGBS samples, either from tissues which have very slight hypomethylation (see Sample 1 in **Fig. 1C**) or tumors with near-complete methylation loss (see Sample 2 in **Fig. 1C**).

We recently used a deep learning neural network approach to establish universal sequence features that are almost entirely predictive of CpG methylation loss or retention in PMD regions of the human genome^32^. We hypothesized that utilizing sequence features associated with DNA methylation loss and exploiting the variation patterns among different CpGs within PMDs could improve the predictive models used in these tools (**Supplementary Fig. 1A-D;** see **Methods)**. To this end, we developed a sequence-aware PMD calling method based on the Hidden Markov Model (HMM) used in MethylSeekR (**Fig. 1C**; see **Methods**), which was termed Multi-model PMD SeekR (MMSeekR). Importantly, using tumor samples from the Blueprint consortium, we showed that MMSeekR outperformed both MethylSeekR and MethPipe (**Supplementary Fig. 1E-F**). Indeed, MMSeekR successfully identified PMD fractions consistently across all samples (using common PMDs as benchmark, top bar, **Supplementary Fig. 1E** and **Supplementary Table 2**). MethylSeekR performed well in general, but was noisier and failed on several samples (**Supplementary Fig. 1E**, red arrows). MethPipe performs poorly on samples with a small degree of PMD methylation loss; indeed, this tool failed to identify PMD in almost half of these samples (**Supplementary Fig. 1E** and **Supplementary Table 2**). PMD has been shown to exhibit cancer type specificity^22,23^, which can also be used to evaluate the performance of these methods. Notably, MMSeekR almost completely separated different cancer types, while both MethylSeekR and MethPipe produced much less clean separation (**Supplementary Fig. 1F**).

Encouraged by these results, we next applied MMSeekR to our esophageal samples. Importantly, Principal Component Analysis (PCA) using PMDs identified by three different methods again confirmed that MMSeekR outperformed MethylSeekR and MethPipe, completely separating EAC and ESCC samples (**Fig. 1D** and **Supplementary Fig. 1G**). Interestingly, nonmalignant samples clustered together with the corresponding cancer subtype. We also provided exemplary PMDs that failed to be identified by either MethPipe (**Fig. 1E**) or MethylSeekR (**Fig. 1F**).

### Characterization of shared and subtype-specific PMDs in esophageal samples

We performed a genome-wide annotation of PMDs on a sample-by-sample basis (**Fig. 2A**). Consistent with our earlier report^19^ and the genome-wide analysis (**Fig. 1B**), PMDs showed a slight decrease of DNA methylation in nonmalignant samples and lost methylation further in tumors. Notably, PMDs exhibited high inter-sample heterogeneity in both their depth (i.e., DNA methylation beta value) and breadth (i.e., genomic location). Indeed, the genome fraction covered by PMDs varied markedly across samples, ranging from 24.3% to 63.4% (**Supplementary Fig. 2C**). We categorized these methylation domains into 4 groups based on the frequencies of their occurrence in our cohort: shared PMDs, EAC-specific PMDs, ESCC-specific PMDs and shared HMDs (**Fig. 2B** and **Supplementary Fig. 2D-E;** also see **Methods**). Interestingly, EAC-specific PMDs covered significantly more of the genome than ESCC-specific PMDs (121.9Mb *vs*. 12.4Mb). To verify our results, we used solo-WCGW CpGs, which lose methylation faster than other CpGs^19^, to measure the average methylation loss within the 4 domain groups. In EAC samples, shared PMDs and EAC-specific PMDs had lower methylation levels than the other two groups, as expected (**Fig. 2C, left panel**). Reciprocally in ESCC samples, shared PMDs and ESCC-specific PMDs had lower methylation levels (**Fig. 2C, right panel**). Independent cohorts from either the TCGA (**Fig. 2D)** or other individual studies (**Supplementary Fig. 2F-G**) further validated these subtype-specific patterns of DNA methylation loss. Since PMDs are associated with the HiC B compartment^17,23^, we next mathematically modeled the A/B chromatin compartments for each esophageal cancer subtype using a method based on the HM450k array^33^. Indeed, subtype-specific PMDs were enriched in B compartments in a subtype-specific manner (**Fig. 2E**). By contrast, shared PMDs showed, as anticipated, no such specificity (**Supplementary Fig. 2H**). PMD regions were also reported to have higher somatic mutation rate compared with non-PMD regions in cancer^34,35^. We analyzed the whole-genome sequencing (WGS) dataset from the OCCAMS (which has the largest number of EAC samples), finding a significantly higher somatic mutation rate in EAC-specific PMDs than in either ESCC-specific PMDs or HMDs (**Fig. 2F, left panel**). A reciprocal pattern was observed in the largest ESCC WGS cohort (**Fig. 2F, right panel**).

**Figure 2.**
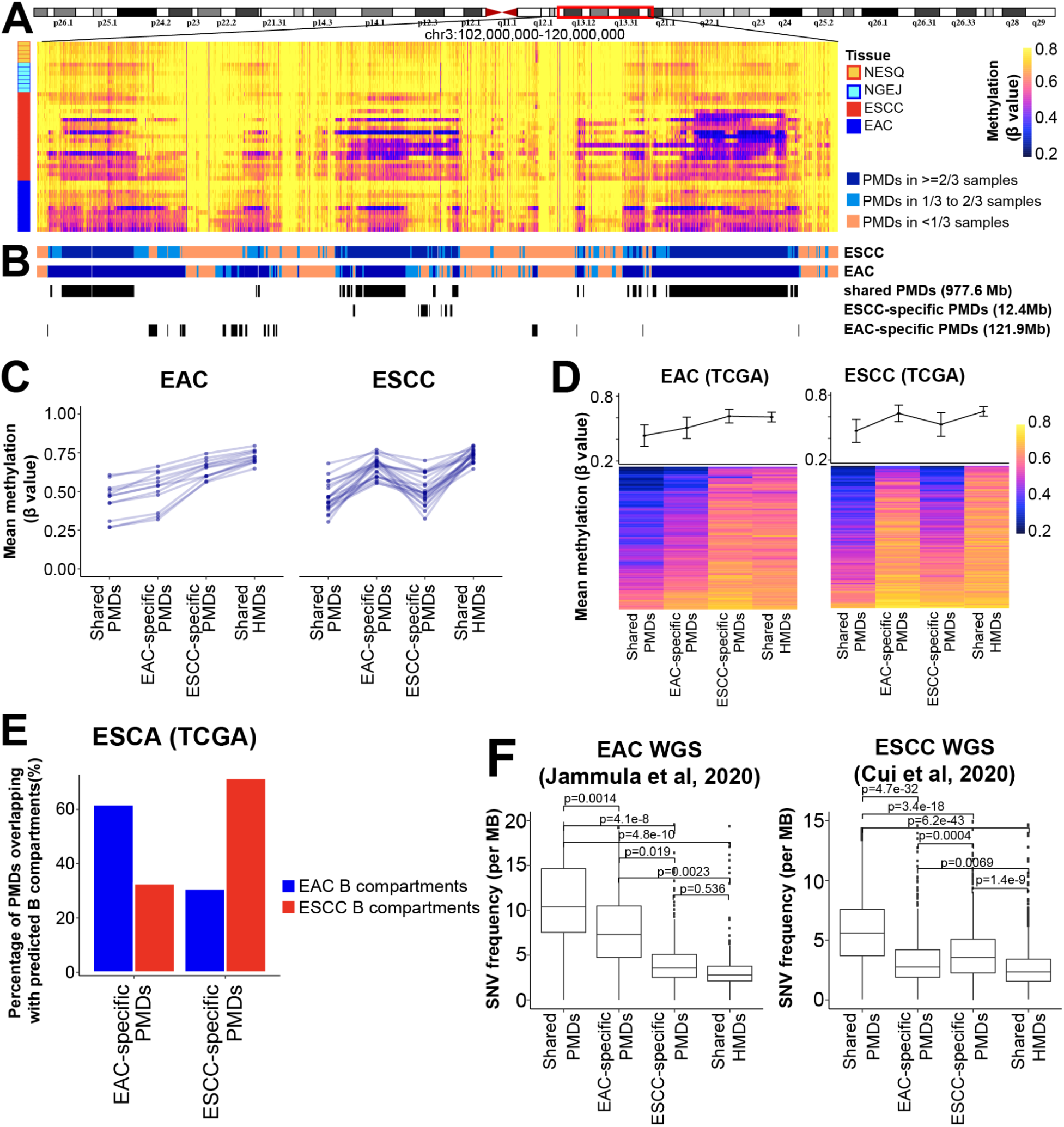
Characterization of shared and subtype-specific PMDs. **(A)** A representative window of DNA methylation profiles from 45 esophageal samples. Average methylation values are shown in consecutive and non-overlapping 10-kb tiles. CGI regions were masked using the annotation from Irizarry et al^76^. **(B)** Different PMD categories were identified based on the frequency and overlap between the two esophageal cancer types. **(C)** Line plots showing average methylation levels for different PMD categories in esophageal tumors, where each line represents one sample. **(D)** Similar line plot patterns were observed using TCGA methylation datasets, showing the mean and standard deviation across samples. Each row in the heatmap below shows an individual sample. **(E)** Bar plots showing the percentage of WGBS PMDs overlapping with chromatin B compartments, which were predicted using TCGA methylation datasets and analyzed by minfi package. **(F)** Somatic mutation rates based on WGS in the indicated studies, calculated separately for each of the WGBS PMD categories.

At the transcription level, PMDs are reported to be less transcriptionally active than HMDs. We confirmed that subtype-specific PMDs were associated with low levels of gene expression specifically in the corresponding subtypes **(Fig. 3A-B)**. To explore the biological implication of subtype-specific PMDs, we performed Cistrome-GO analysis using genes which were under-expressed in the subtype-specific PMD regions, finding that biological processes characteristic for the other subtype were enriched and repressed **(Fig. 3C-D)**. Specifically, pathways of cornification, keratinocyte differentiation and epidermis development, which are central to squamous cell differentiation and function, were enriched and inactive in EAC-specific PMDs **(Fig. 3C)**. For example, many keratinocyte-specific genes were clustered within EAC-specific PMDs (**Fig. 3E**, **left panel**) and downregulated in EAC tumors (**Fig. 3F**, **upper panel**). On the other hand, pathways important for gastrointestinal cell function, such as digestive system process, intestinal absorption, lipid metabolic process and O-glycan processing, were enriched and suppressed in ESCC-specific PMDs (**Fig. 3D**). The right panel of **Fig. 3E** shows as an example that SLC2A2, which contributes to digestive system process and absorption, was located in ESCC-specific PMDs and downregulated in ESCC samples (**Fig. 3F**, **lower panel**). These results suggest that subtype-specific PMDs contain inactive genes which are associated with cell-type-specific functions.

**Figure 3.**
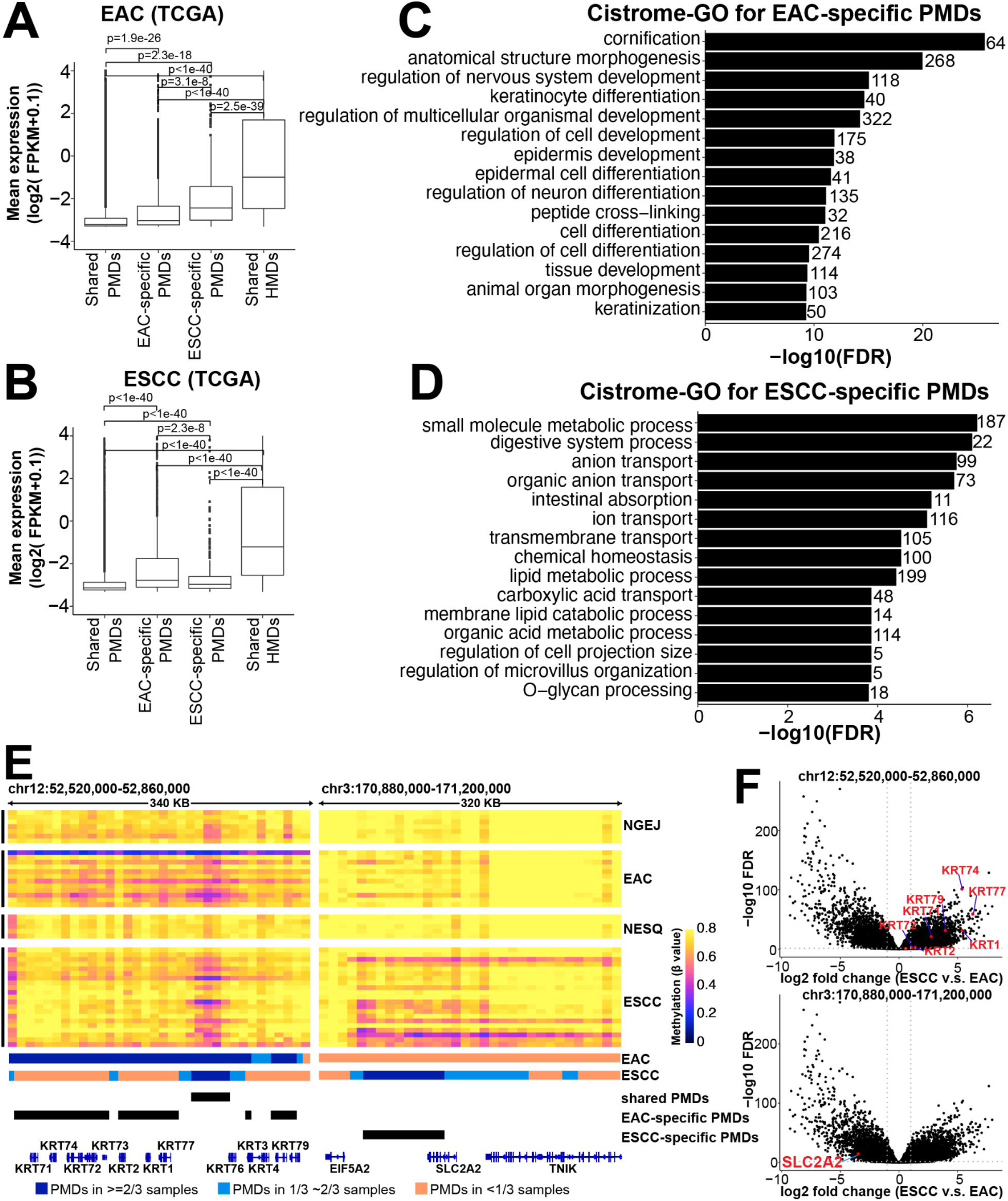
Subtype-specific PMDs control cell-type-specific functions. **(A-B)** In both EAC **(A)** and ESCC **(B)**, genes covered by PMDs are expressed at lower levels than those in non-PMDs in a cancer-specific manner. **(C-D)** Cistrome-GO enrichment analyses using either EAC-specific **(C)** or ESCC-specific **(D)** PMDs and the downregulated genes within them. The top 15 most significant pathways are shown, and the number of genes enriched in each pathway is shown on the right. **(E)** Two representative genome windows showing the methylation profiles of EAC-specific (left panel) and ESCC-specific PMDs (right panel). CGI regions were masked using the annotation from Irizarry et al^76^. **(F)** Volcano plots showing that genes residing within genome domains in **(E)** are downregulated in corresponding cancer subtypes.

### H3K36me2 is inversely associated with PMDs in a cell-type-specific manner

Both H3K36me2 and H3K36me3 were observed to recruit DNA methyltransferases (DNMT3A^36^ and DNMT3B^37^, respectively) to maintain DNA methylation levels in large chromatin domains. H3K36me3 is enriched in gene bodies of active transcripts, while H3K36me2 covers larger multi-gene domains. Indeed, we have previously shown that the deposition of H3K36me3 is inversely associated with PMD distribution^19^. Here, we further hypothesized that H3K36me2 also contributed to maintaining DNA methylation levels, and the histone modification by this mark might affect the genomic distribution of PMDs and HMDs. To test this, we performed H3K36me2 ChIP-seq in both EAC and ESCC cell lines. Indeed, shared HMDs (black line) showed high H3K36me2 intensity in both cell types, while shared PMDs (purple line) exhibited the lowest signals (**Fig. 4A**). EAC-specific PMDs (blue line) had low H3K36me2 levels in EAC cells but high H3K36me2 levels in ESCC cells. The reciprocal pattern was observed in ESCC-specific PMDs (red line). For example, H3K36me2 signals were undetectable in an EAC-specific PMD covering the loci of *XR_945002.2* and *XR_945004.2* in EAC cells, but were strong in ESCC (**Fig. 4B, right panel**). On the other hand, shared HMDs such as the one covering the *VSP8* gene were decorated highly with H3K36me2 in both cell types (**Fig. 4B, left panel**).

**Figure 4.**
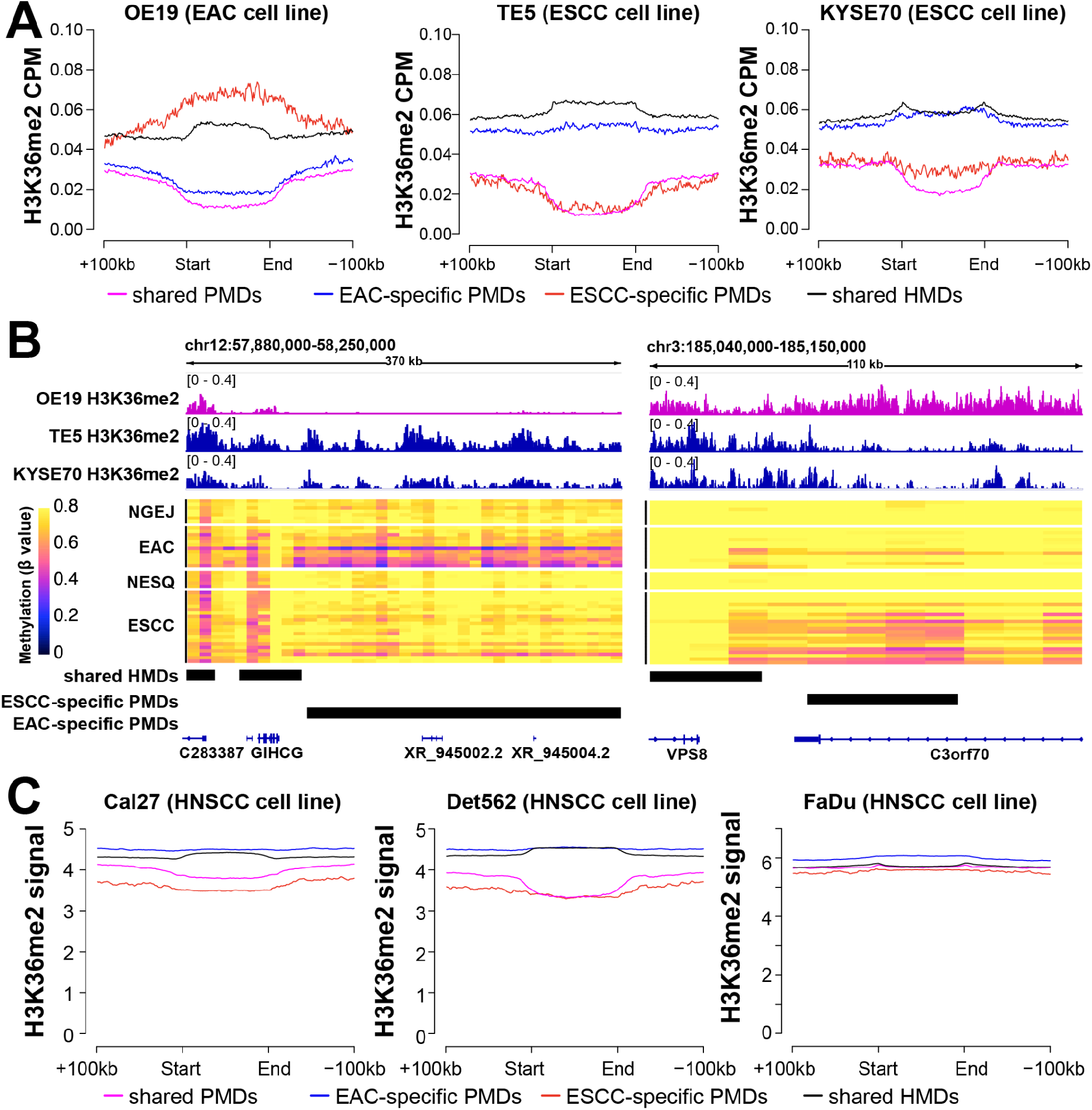
The H3K36me2 mark is inversely associated with PMDs in a cell-type-specific manner. **(A)** Aggregation plots of H3K36me2 ChIP-seq levels in esophageal cancer cell lines across four different PMD categories: shared PMDs, EAC-specific PMDs, ESCC-specific PMDs, shared HMDs. **(B)** Representative genomic loci showing H3K36me2 signal from ChIP-seq, and subtype-specific PMDs from WGBS data. CGI regions were masked using the annotation from Irizarry et al^76^. **(C)** Aggregation plots of H3K36me2 ChIP-seq levels in HNSCC cell lines across four different PMD categories. Bigwig files of the H3K36me2 ChIP-seq signal were obtained from GSE149670.

To further verify these results, we interrogated public H3K36me2 ChIP-seq data from HNSCC cell lines (squamous cancer highly similar to ESCC in terms of cell-of-origin and epigenome). Indeed, a similar pattern of H3K36me2 distribution to ESCC was observed in Cal27 and Det562 HNSCC cells. Specifically, both shared PMDs and ESCC-specific PMDs harbored low signals in HNSCC cell lines, while high H3K36me2 levels were found in HMDs and EAC-specific PMDs (**Fig. 4C**). However, FaDu appeared to be an outlier, showing invariably high levels across different regions (**Fig. 4C**), which warrants further investigation. Together, these results demonstrate a prominent depletion of H3K36me2 mark in PMDs in a cell-type-specific manner, which is likely owing to the finding that H3K36me2 promotes the maintenance of DNA methylation by recruiting DNMT3A.

### Subtype-specific differentially methylated regions (DMRs) in esophageal cancer

We next sought to investigate differentially methylated regions (DMRs) at small genomic scales, given their direct roles in transcriptional regulation. However, our above results suggest an overwhelming, global effect of PMD hypomethylation in tumor samples, which can strongly affect the calling of focal DMRs. Indeed, PCA analysis of the most variable CpGs genome-wide revealed that PC1, the most significant component, was clearly driven by methylation loss at PMDs (**Supplementary Fig. 3A**).

To factor out the effect of PMD hypomethylation, we masked any PMD found within two thirds of either EAC or ESCC samples (**Supplementary Fig. 3B**). We re-performed the PCA analysis, finding that the two cancer subtypes were completely separated by PC1, which was the most significant component and accounted for 42.2% of the total methylation variance (**Supplementary Fig. 3C, left panel**). In addition, nonmalignant and tumor samples were separated along PC2, and all NESQ samples were clustered closely together despite being generated from two different cohorts. Notaly, this approach removed most correlation with the global methylation level (**Supplementary Fig. 3C, right panel**). Thus, it is critical to remove the effects of global hypomethylation when investigating cancer-associated methylation features outside PMDs.

We next identified DMRs between EAC and ESCC samples within the PMD-subtracted genome described above (~46.5% of the genome). Under the cutoff of q value < 0.05 and absolute delta methylation change > 0.2, a total of 7,734 DMRs were hypomethylated in EAC and 5,470 in ESCC (**Fig. 5A**). As expected, hypomethylated DMRs (hypoDMRs) had low average methylation levels in corresponding subtypes (**Supplementary Fig. 3D-E**). The majority of DMRs were about 1-2 kb long and located mostly in intronic and intergenic regions (**Fig. 5B**), similar to that of the random background (**Supplementary Fig. 3F**). To investigate the epigenomic characteristics of hypoDMRs, we systematically evaluated the chromatin accessibility at these regions, using the ATAC-seq data from the TCGA^38^ and H3K27ac ChIP-seq data from previous studies^39–42^. Relative to random background regions, EAC hypoDMRs were accessible exclusively in EAC samples, and ESCC hypoDMRs exclusively in ESCC samples (**Fig. 5C-D**). Additionally, EAC hypoDMRs had high H3K27ac signals in 70% (5/7) of EAC cell lines (**Supplementary Fig. 3G**). A similar observation was made in ESCC cell lines (**Supplementary Fig. 3H**). These data demonstrate that hypoDMR regions are associated with accessible chromatin and active histone marks.

**Figure 5.**
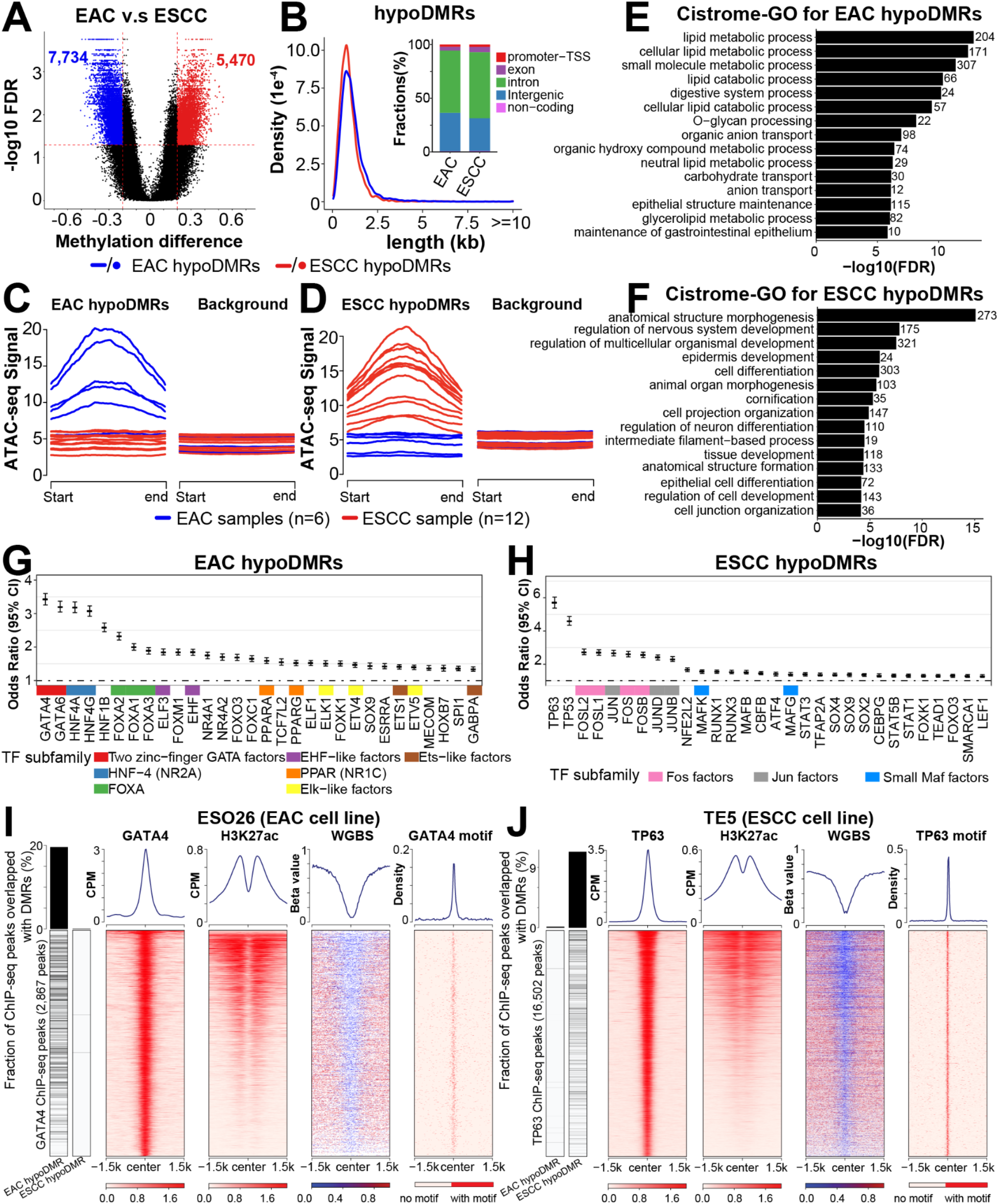
Subtype-specific DMRs in esophageal cancer. **(A)** Cancer hypoDMRs were identified from the comparison between EAC and ESCC tumors. Regions with FDR < 0.05 and absolute delta methylation levels > 0.2 were identified as DMRs. **(B)** Density plots showing the size distribution of hypoDMRs; stacked bar plots displaying fractions of hypoDMRs that overlap with different genomic features. **(C-D)** Aggregation plots of ATAC-seq signals from esophageal cancer samples within EAC **(C)** or ESCC **(D)** hypoDMRs or random genomic regions (background), which contained 10-times randomly selected regions with the same CpG density. ATAC-seq signals were obtained from the TCGA and normalized with the CPM method. **(E-F)** Cistrome-GO enrichment analyses using EAC **(E)** or ESCC **(F)** hypoDMRs and upregulated genes in the corresponding subtype. Top 15 most significant pathways are shown. The number of genes enriched in each pathway is shown on the right. **(G-H)** Transcription-factor-binding motif sequences were identified by the ELMER^77^ method using EAC **(G)** or ESCC **(H)** hypoDMRs as the foreground and random regions as the background. **(I-J)** The most strongly enriched TFs in EAC (GATA4) **(I)** and ESCC (TP63) **(J)** were chosen for the experimental validation, using TF ChIP-seq, H3K27ac ChIP-seq and WGBS in matched cell lines. Peaks overlapping with subtype hypoDMRs are shown on the left; the percentages of overlapped peaks are expressed in the column plots.

To explore the relevance of DMRs in gene transcription, we assigned each hypoDMR to the closest genes annotated by HOMER^43,44^, and performed correlational analyses using TCGA transcriptomic data of esophageal cancers. Consistent with prior findings^43^, about 30% (3,986/13,204) of the DMRs were associated with differentially expressed genes **(Supplementary Fig. 3I)**. Expectedly, an inverse correlation between DNA methylation and gene expression accounted for the majority (~59%) of these associations, and these DMRs had a larger overlap with promoter and enhancer regions **(Supplementary Fig. 3J)**. Importantly, functional annotation using the Cistrome-GO method revealed that subtype hypoDMRs were enriched in cell-type-specific biological processes. For example, lipid metabolic process, digestive system process and O-glycan processing, which are housekeeping functions for gastrointestinal columnar cells, were specifically enriched in EAC hypoDMRs **(Fig. 5E)**. On the other hand, epidermis development, cornification and epithelial cell differentiation, which are unique to squamous cells, were enriched in ESCC hypoDMRs **(Fig. 5F)**. These results indicate that a large number of hypoDMRs regulate the transcription of cell-type-specific genes.

We next performed sequence motif enrichment analysis of hypoDMRs, which have previously been associated with transcription-factor-binding sites^17,22,45^. A number of known esophageal cell-specific transcription factors were identified, including GATA4/6, HNF4A/G, HNF1B, ELF3, EHF in EAC^39,46,47^ and TP63, SOX2 and MAFB in ESCC^41,48^ (**Fig. 5G-H**). To validate these results, we focused on the top-ranking transcription factors (GATA4 for EAC, TP63 for ESCC). Specifically, we performed WGBS in an EAC cell line (ESO26) where we previously generated ChIP-seq data for GATA4 and H3K27ac. Indeed, GATA4 ChIP-seq peaks were associated with high H3K27ac signal, DNA hypomethylation and GATA4 binding motif sequence (**Fig. 5I)**. Moreover, ~20% of GATA4 peaks overlapped with EAC hypoDMRs. In sharp contrast, almost no GATA4 peaks were found in ESCC hypoDMRs (**Fig. 5I, left bars**). We similarly performed WGBS on an ESCC cell line (TE5), and analyzed TP63 ChIP-Seq data that we generated in the same sample. We noted consistent patterns and significant overlap with ESCC hypoDMRs in this ESCC-specific transcription factor, and almost no overlap with EAC hypoDMRs (**Fig. 5J**). These results demonstrate that subtype-specific DMRs are occupied by cell-type-specific transcription factors and contribute to regulation of cell-type-specific functions.

### Identification of tumor-specific hypoDMRs

To identify tumor-specific hypoDMRs from the above subtype-specific DMRs and to investigate their role in cancer biology, we next performed a methylation comparison between tumors and their corresponding nonmalignant samples for each hypoDMR. We found that 25.5% (1,972/7,734) of EAC hypoDMRs (**Fig. 6A**) and 12.0% (654/5,470) of ESCC hypoDMRs (**Supplementary Fig. 4A**) had significantly lower (FDR<0.05) methylation levels in tumors than corresponding nonmalignant samples, which were referred to as “tumor specific hypoDMRs (ts-hypoDMRs)”, while the rest were referred to as “cell-type-specific DMRs (cts-hypoDMRs)”. Ts-hypoDRMs were distributed in both intergenic and intronic domains, similar to hypoDMRs overall and the random background (**Fig. 6B** and **Supplementary Fig. 4B**). Between 18.0-21.4% of ts-hypoDMRs were correlated with the expression of nearest genes (**Supplementary Fig. 4C-D**). Importantly, ts-hypoDMRs were strongly enriched in cancer-related pathways such as cell cycle progression (in both EAC and ESCC), and extracellular structure organization in ESCC (**Fig. 6C-D)**. These data suggest that ts-hypoDMRs are associated with genes which contribute to tumorspecific functions.

**Figure 6.**
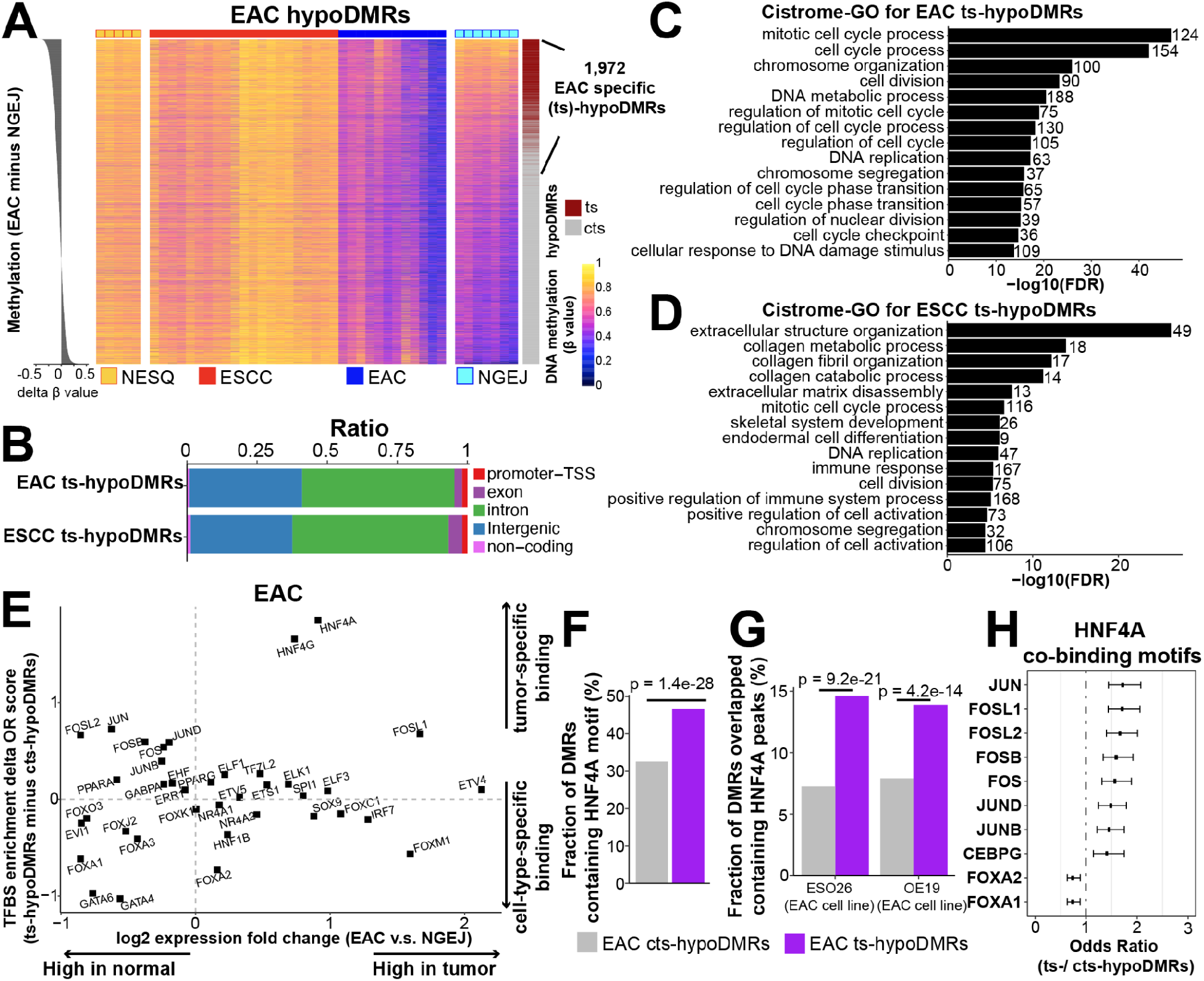
Identification of tumor-specific hypoDMRs. **(A)** Heatmaps showing DNA methylation levels for each EAC hypoDMR. Each column denotes one sample and the row was ordered by the delta mean methylation between EAC and NGEJ (left). EAC ts-hypoDMRs were identified using a one-tailed t test between EAC tumor and NGEJ samples (right) with the FDR cutoff < 0.05. **(B)** Stacked bar plots showing fractions of ts-hypoDMRs that overlap with different genomic features. **(C-D)** Cistrome-GO enrichment analyses using either EAC **(C)** or ESCC **(D)** ts-hypoDMRs and the upregulated genes in each subtype compared with corresponding nonmalignant samples. Top 15 most significant pathways are shown. **(E)** Scatter plots showing transcription-factor-binding sites that were enriched in EAC ts-hypoDMRs compared with cts-hypoDMRs. The X axis represents the expression fold change between EAC and matched nonmalignant GEJ samples. The Y axis shows the delta enrichment score of transcription-factor-binding sites between EAC ts- and cts-hypoDMRs. Expression data were from the TCGA and motif enrichment analyses were performed by the ELMER method. **(F)** EAC ts-hypoDMRs contained significantly more HNF4A-recognition motifs compared with cts-hypoDMRs. **(G)** More HNF4A peaks overlapped with ts-hypoDMRs than cts-hypoDMRs. Peaks were called from HNF4A ChIP-seq in ESO26 and OE19 cell lines. **(H)** HNF4A was predicted to co-occupy with the AP-1 family in ts-hypoDMRs, while with FOXA1/2 in cts-hypoDMRs. Sequence motif analysis was performed using ts-*vs*. cts-hypoDMRs containing HNF4A motifs. Significant transcription factors with FDR < 0.05 are shown. OR value over 1 represents higher enrichment in ts-hypoDMRs, while below 1 represents higher enrichment in cts-hypoDMRs.

The identification of ts-hypoDMRs and cts-hypoDMRs allowed us to further investigate properties of tumor-specific regulatory regions *vs*. cell-type-specific regulatory regions. This is particularly helpful for the epigenetic understanding of ESCC and EAC, which contain both tumor- and cell-type-specific features. In addition, lineage-specific developmental factors have been shown to promote malignant cell states^49,50^, and thus it is important to distinguish their functional contribution to normal development *vs*. cancer biology. To this end, we performed motif enrichment analysis to identify transcription-factor-binding sites that were unique to either ts- or cts-hypoDMRs, and integrated expression patterns of the corresponding transcription factors. For EAC, this approach revealed cancer-upregulated transcription factors which favored binding ts-hypoDMRs, including HNF4A, HNF4G, and FOSL1 (upper right corner of **Fig. 6E**). In comparison, the lower left corner of **Fig. 6E** contained cancer-downregulated transcription factors which preferred occupying cts-hypoDMRs, including GATA4/6 and FOXA, which are well-recognized for their key roles in the development of gastrointestinal cell lineage^51,52^. The top factor for ts-hypoDMR, HNF4A, had its binding motif in 46.6% ts-hypoDMRs but only 32.6% cts-hypoDMRs (**Fig. 6F**). Indeed, ChIP-seq data of HNF4A in EAC cell lines (ESO26 and OE19) validated this bias: HNF4A binding peaks overlapped with 14.2% ts-hypoDMRs but only 7.6% cts-hypoDMRs (**Fig. 6G**). To identify factors that may facilitate recruitment of HNF4A specifically to hypoDMRs, we performed enrichment analyses restricted within HNF4A-motif-containing hypoDMRs. Interestingly, AP-1 motifs (such as JUN, FOSL1, FOSL2 and FOSB) were enriched in these HNF4A^+^ ts-hypoDMRs, while FOXA1/2 in cts-hypoDMRs (**Fig. 6H**). A parallel analysis was performed in ESCC, which identified a number of tumor-specific factors, including RUNX1/3, SOX2/4 and CEBPA/B (**Supplementary Fig. 4E**). This distinct pattern of co-occurring motifs between ts- and cts-hypoDMRs in EAC is noteworthy, considering that AP-1 family transcription factors contribute to EAC tumor development^53^ while FOXA1/2 are required for normal gastrointestinal cell development^52^. It is also notable that our analysis identified FOSL1 as an AP-1 factor due to its high tumor expression **(Fig. 6E)**.

### PMDs and hypoDMRs exhibit strong cell-type-specific epigenomic features

The above data identified both cell-type- and cancer-specific methylation differences in tumor hypoDMRs, and we next asked whether tumor PMDs likewise harbor both of these two types of methylation differences. In subtype-specific PMDs that were defined based on tumor methylomes alone, nonmalignant tissues notably exhibited the same pattern of methylation changes as their malignant counterparts (**Fig. 7A**). For example, EAC-specific PMDs had low methylation levels in NGEJ but high in NESQ **(Fig. 7A, left)**, and a reciprocal pattern was found in ESCC-specific PMDs **(Fig. 7A, right)**. Statistically, a large subset of subtype-specific PMDs (33.0% for EAC and 26.5% for ESCC) were already hypomethylated in their respective nonmalignant samples (**Fig. 7B**). The same analyses for hypoDMRs confirmed that more than 80% of subtype hypoDMRs significantly decreased DNA methylation in their corresponding nonmalignant samples (**Fig. 7C-D**). These data demonstrate that a substantial fraction of both subtype-specific PMDs and hypoDMRs identified from tumor samples reflect methylation differences present in normal counterparts. Nonetheless, while the genomic locations of PMDs are established in normal samples, the degree of methylation loss is significantly higher in tumors (**Fig. 2C** and **Supplementary Fig. 3D**).

**Figure 7.**
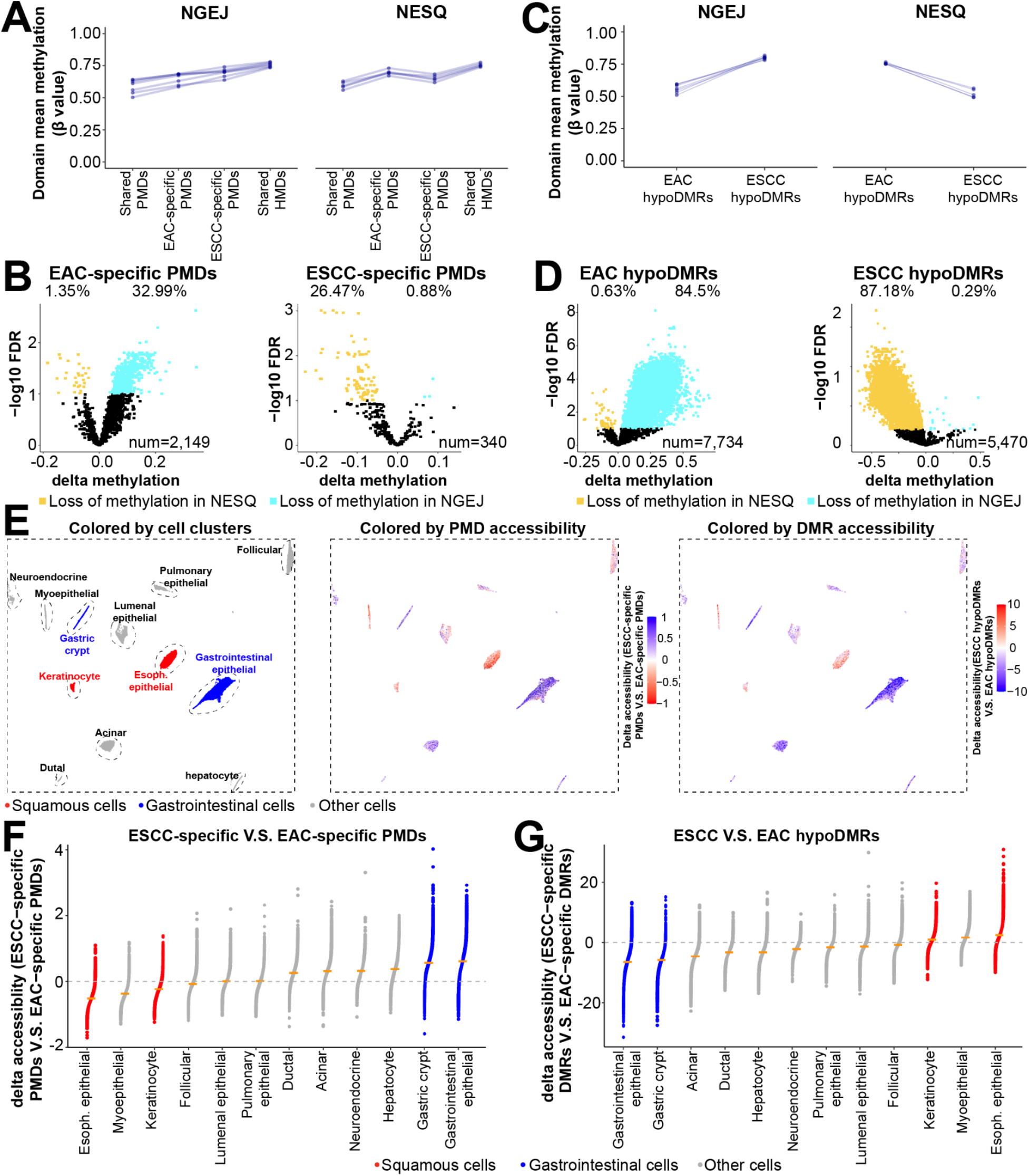
PMDs and hypoDMRs exhibit strong cell-type-specific epigenomic features. **(A)** Line plots showing average methylation levels for different PMD or **(C)** hypoDMR categories comparing two types of nonmalignant esophageal samples; these changes in nonmalignant samples are similar to those seen in tumors **(Fig. 2C, Supplementary Fig. 3D-E)**. **(B)** Volcano plots showing average methylation levels for different PMD or **(D)** hypoDMR categories in nonmalignant esophageal samples. Regions with significant differences were determined by two-tailed t test with the FDR cutoff < 0.1. **(E)** UMAP plots showing cell clusters (left), ATAC-seq levels in ESCC-*vs*. EAC-specific PMDs (middle) or in ESCC-*vs*. EAC-specific hypoDMRs (right). Single-cell ATAC-seq values and the cluster scheme were from Zhang et al. Total cell number is 146,305. **(F-G)** Dot plots showing, at the sample level, delta ATAC-seq values in ESCC-*vs*. EAC-specific PMDs **(F)** or in ESCC-*vs*. EAC-specific hypoDMRs **(G)**.

To understand further PMDs and hypoDMRs in normal samples, we analyzed public single-cell ATAC-seq data from 146,305 normal epithelial cells across 24 tissues (including esophageal samples)^54^, by measuring the chromatin accessibility of our subtype-specific PMDs or hypoDMRs. This is premised on the fact that focal ATAC-seq peaks are almost always DNA demethylated^38^, and reduced ATAC-seq signals measured in large genomic windows reflect the Hi-C B compartment which results in PMD hypomethylation^17,23^. The published single-cell unsupervised clustering contains a cluster of esophageal squamous epithelial cells (red dots in **Fig. 7E, left panel**), the recognized cell-of-origin for ESCC. With respect to EAC, although its cell-of-origin is still under intense investigation, the epigenome is likely close to gastrointestinal epithelial cells (blue dots **Fig. 7E, left panel**). Importantly, normal esophageal squamous cells showed the lowest chromatin accessibility in ESCC-specific PMDs; reciprocally, normal gastrointestinal epithelial cells had the lowest ATAC-Seq signals in EAC-specific PMDs (**Fig. 7E, middle panel; quantified in Fig. 7F**). In addition, keratinocytes, which belong to squamous cell type, also had low ATAC-Seq signals in ESCC-specific PMDs. In sharp contrast to subtype-specific PMDs, no difference was observed in either shared PMDs or HMDs in this single-cell analysis (**Supplementary Fig. 5C**). We performed the same analysis for hypoDMRs, finding that ESCC hypoDMRs had the highest accessibility in squamous cells while EAC hypoDMRs were more open in gastrointestinal epithelial cells (**Fig. 7E, right panel; quantified in Fig. 7G**). These single-cell results confirmed that both PMDs and hypoDMRs have strong normal cell-type-specificity.

### Pan-cancer analysis of subtype-specific PMDs and hypoDMRs

The above results also suggest that PMDs and hypoDMRs that we identified in ESCC and EAC may be shared with other squamous and gastrointestinal adenocarcinomas, respectively. To test this, we analyzed TCGA pan-cancer samples, since the TCGA multi-omic clustering scheme^55^ has identified the pan-gastrointestinal cluster (adenocarcinomas from esophagus, stomach and colon, blue samples in **Fig. 8A**) and the pan-squamous cluster (squamous cancers from esophagus, head and neck, lung, cervix and bladder, red samples in **Fig. 8A**). We first measured the methylation changes between subtype-specific PMDs and hypoDMRs across all 33 cancer types (**Fig. 8B-E)**. Importantly, most pan-gastrointestinal tumors lost DNA methylation in EAC-specific PMDs, while most pan-squamous tumors had reduced methylation in ESCC-specific PMDs (**Fig. 8B and 8D**). Highly consistent results were observed in subtype hypoDMRs (**Fig. 8C and 8E**). In contrast, no specific pattern was found in shared PMDs and HMDs (**Supplementary Fig. 5D**), as anticipated.

**Figure 8.**
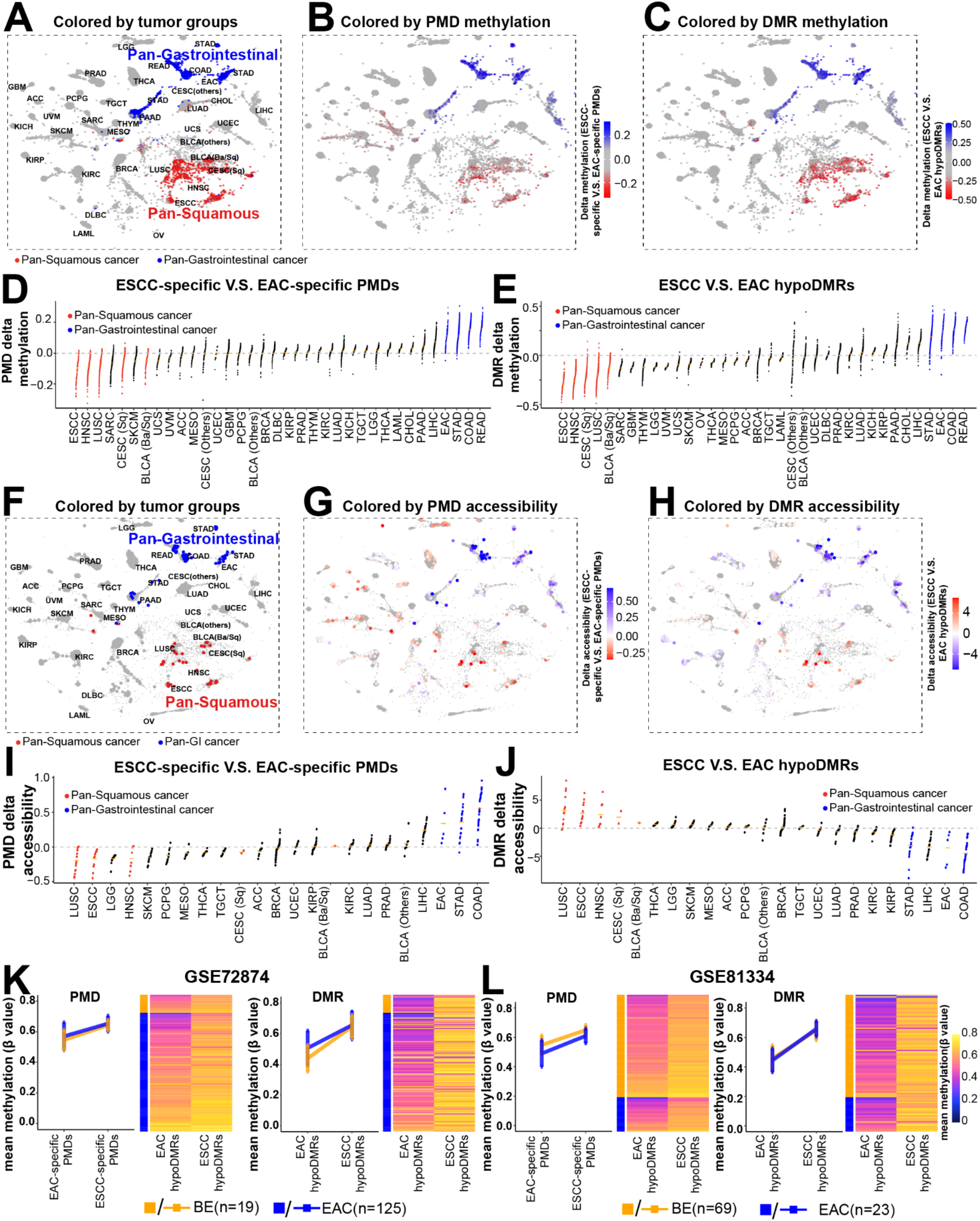
Analyses of PMDs and hypoDMRs in pan-cancer datasets. **(A-C)** TCGA tumormap showing cancer type clusters **(A)**, DNA methylation levels in ESCC-*vs*. EAC-specific PMDs **(B)**, or in ESCC-*vs*. EAC-specific hypoDMRs **(C)**. DNA methylation data were obtained from the TCGA project. The TCGA-based clustering scheme denotes Pan-Gastrointestinal cancers (COAD, READ, STAD and EAC) and Pan-squamous cancers (ESCC, HNSC, LUSC and a subset of CESC and BLCA) are shown **(A)**. The number of samples is 8,915. The detailed study name of TCGA Study Abbreviations are listed in https://gdc.cancer.gov/resources-tcga-users/tcga-code-tables/tcga-study-abbreviations **(D)** and **(E)** Dot plots quantification of the methylation differences in **(B)** and **(C)**, respectively. **(F)** t-SNE plots showing cancer type clusters, **(G)** ATAC-seq levels in ESCC-*vs*. EAC-specific PMDs or in **(H)** ESCC- *vs*. EAC-specific hypoDMRs across tumor samples. ATAC-seq data were downloaded from the TCGA project. The number of samples is 362. **(I)** and **(J)** Dot plots quantification of the ATAC-seq values in **(G)** and **(H)**, respectively. **(K-L)** Line plots and heatmaps respectively showing average and individual methylation levels in BE and EAC samples from two different public datasets.

We next analyzed the ATAC-seq data, which is available from a small subset of TCGA bulk tumors^38^, shown based on multi-omic clustering from ref^55^ in **Fig 8F**. Importantly, consistent with the single-cell ATAC-Seq results from healthy tissues, pan-squamous cancers showed the lowest chromatin accessibility in ESCC-specific PMDs and highest accessibility in ESCC hypoDMRs, and the reciprocal results were obtained in pan-gastrointestinal cancers (**Fig. 8G-J**). Again, as negative controls, shared PMDs and HMDs failed to generate this distinguishing epigenetic pattern (**Supplementary Fig. 5E**).

These results prompted us to further investigate premalignant lesions, with the hypothesis that these methylation changes are pre-established in normal cells and preserved during the onset of neoplastic transformation. To address this, we interrogated public methylation data on BE, a recognized precursor to EAC, from two different studies^7,8^. Importantly, the methylation patterns of BE samples were highly comparable with EAC tumors, showing reduced methylation levels in both EAC-specific PMDs and hypoDMRs in two different cohorts (**Fig. 8K-L**). Overall, these data strongly suggest that epigenomic changes of PMDs and hypoDMRs occur in normal cells and are maintained in cancer, which further loses methylation in PMDs and gains additional DMRs. Moreover, these region-specific epigenomic regulations are shared across related cell types.

## Discussion

We generated one of the largest WGBS datasets in esophageal cancer to date, and here we focused on the analyses of PMDs (large scale) and DMRs (small scale) and revealed novel epigenomic properties of these regions. PMDs are megabase-long genomic regions with decreased DNA methylation, coinciding with heterochromatic late-replicating domains and Hi-C B domains^17^. PMDs reflect long-range chromatin organization that help orchestrate gene expression programs and can influence replication timing and 3D genome organization^24,33,56–58^. In addition, PMDs are associated with increased genomic instability and possibly activation of transposable elements (TEs)^19,21^. Nevertheless, apart from these correlational observations, we have only limited mechanistic understanding of the origin and regulation of cancer PMD. Moreover, direct mechanisms linking PMDs to gene transcription remain to be established. Thus, a deeper characterization of PMD is warranted, which first requires an accurate and sensitive identification of these large domains from WGBS data. However, current PMD callers, including MethylSeekR and MethPipe, either are insensitive for the identification of shallow PMDs, or fail to call PMDs in tumor samples with extreme hypomethylation.

We have previously demonstrated that a local sequence context (solo-WCGW) is a strong determinant of DNA methylation loss at CpGs^19^. Extending this finding, we recently performed deep learning using the neural network method, and established universal sequence context features influencing the hypomethylation of CpGs across the genome^32^. Here, we integrated this sequence code into the MethylSeekR program and developed a novel multi-model PMD caller, MMSeekR. Using both the Blueprint tumor WGBS dataset and our esophageal samples, we demonstrated a superior performance of MMSeekR over other current tools. In order to facilitate methodological development in the field of methylome investigation, we have made MMSeekR available at Github as a free software package (https://github.com/yuanzi2/MMSeekR).

The degree of variation of PMD methylation levels (depth) and genomic distribution (breadth) between cancer types was hitherto unclear. Here we observed strong heterogeneity at the PMD methylation level across cancer samples, while nonmalignant samples harbored expectedly shallow PMDs. Moreover, the genome fraction covered by PMDs varied profoundly among different samples, ranging from 24.3% to 63.4%. We identified and characterized subtype-specific PMDs, finding that they were associated with repressive transcription, B compartments and high somatic mutation rate. We previously identified replication timing as a key determinant for methylation loss in PMDs^19^. However, this does not account for the variation in PMD genomic distribution across cell types. By investigation of the genome-wide occupancy of H3K36me2 in different cell types, we noted that H3K36me2 deposition correlated positively with HMD localization, while negatively with PMD in a cell-type-specific manner. Considering that H3K36me2 is able to recruit DNMT3A to maintain the level of DNA methylation^36^, these results suggest that cell-type-specific deposition of H3K36me2 mark facilitates the maintenance of DNA methylation, thereby dictating the genomic distribution of HMDs and PMDs.

At a smaller genomic scale, we identified over ten thousand hypoDMRs between the two subtypes of esophageal cancer. Utilizing their matched nonmalignant samples, we further defined cell-type-*vs*. cancer-specific hypoDMRs. Using motif sequence analysis combined with ChIP-seq, we identified and validated candidate upstream regulators associated with either cell-type- or cancer-specific hypoDMRs. This approach is important for understanding of the transcriptional regulation during tumor development, particularly because increasing evidence has shown that tumor-driving transcription factors are often lineage-specific developmental regulators functionally co-opted to promote malignant cellular states^49,50^. For example, our top candidate, HNF4A, is essential for the epithelial differentiation of the gastrointestinal tract. Consistently, we found that a substantial subset of cell-type-specific hypoDMRs contained HNF4A-binding sequence; these HNF4A^+^ cell-type-specific hypoDMRs were also co-enriched for transcript factors indispensable for normal gut development, such as FOXA1 (**Fig. 6H**). Importantly, compared with cell-type-specific hypoDMRs, HNF4A-binding sequence was significantly more enriched in tumor-specific hypoDMRs (**Fig. 6H)**. Moreover, instead of FOXA1, these HNF4A^+^ tumor-specific hypoDMRs were co-enriched for AP-1 factors, which are well-recognized for their function in promoting EAC malignancy^53^, similar to HNF4A itself^46,47^. Consistently, one of the AP1 factors, FOSL1, has highly enriched binding sites in tumor-specific hypoDMRs as well as upregulated mRNA expression in EAC tumors relative to NGEJ. Together, careful dissection of cell-type- and cancer-specific hypoDMRs suggest that lineage master regulators control both normal and tumor cell transcriptomes, likely by occupying different genomic regions through cooperating with different transcriptional factor partners.

We further characterized the cell-type-specificity of PMDs and DMRs in normal cells. Starting from esophageal samples, we found that a large fraction of methylation changes in both PMDs and DMRs were already evident in normal samples. Pan-tissue single-cell ATAC-seq with 145,594 normal epithelial cells further showed that both PMDs and DMRs identified in esophageal cancer had strong specificity that was evident in related cell types. This was also observed in pan-cancer analyses of both methylation and ATAC-seq data from primary tumors, wherein cancers originating from related cell types exhibited similar profiles of both PMDs and DMRs. Moreover, by measuring cancer precursor lesions, we demonstrated that epigenomic changes of PMDs and DMRs were preserved during the onset of neoplastic transformation. Nonetheless, PMDs in normal samples were much shallower than tumors (**Fig. 2A** and **Fig 2C *vs*. Fig.7A**). Overall, these data highlight the presence of cell-type-specific PMDs and DMRs in normal cell types, which are preserved in malignant cells. To our knowledge, this is the first demonstration of the prominent cell-type-specificity of PMDs across normal, precursor and malignant states. While prior studies have revealed that DMRs contain tissue-specific regulatory regions, here we present a paradigm for distinguishing cell-type-*vs*. cancer-specific regions, and use those to identify tumor-specific regulatory mechanisms.

## Methods

### Cell culture

Esophageal cancer cell lines, TE5, KYSE70, OE19 and ESO26, were grown in RPMI-1640 medium (Gibco, USA), supplemented with 10% FBS (Omega Scientific, USA) and 1% penicillin-streptomycin (Thermo Scientific, USA). All cultures were maintained in a 37 °C incubator supplemented with 5% CO2.

### Whole genome bisulfite sequencing (WGBS)

WGBS of ESO26 or TE5 cells was performed at Novogene, Inc. Briefly, after DNA extraction and quality control (QC), 3 ug DNA of ESO26 or TE5 cells spiked with 26 ng lambda DNA were fragmented by sonication. The sonicated DNA was ligated with different cytosine-methylated molecular barcodes. Next, bisulfite conversion was performed using EZ DNA Methylation-GoldTM Kit (Zymo Research). PCR amplification with KAPA HiFi HotStart Uracil+Ready Mix (Kapa Biosystems) was then applied to the DNA fragments. The clustering of index-coded DNA samples were sequenced using the Illumina Hiseq 2500 platform.

### H3K36me2 chromatin immunoprecipitation sequencing (ChIP-Seq)

Ten million esophageal cancer cells were harvested and transferred into 15 ml tubes, followed by fixing with 4 ml of 1% paraformaldehyde for 10 min under room temperature. The reaction was stopped by 2 ml of 250 mM of glycine. Cell samples were rinsed twice by 1X PBS and lysed by 1 ml of 1X lysis/wash buffer (150 mM NaCl, 0.5 M EDTA pH 7.5, 1M Tris pH 7.5, 0.5% NP-40). Cell pellets were next resuspended using shearing buffer (1% SDS, 10 mM EDTA pH 8.0, 50 nM Tris pH 8.0) followed by sonication using a Covaris sonicator. Subsequently, debris was removed by centrifuge and supernatants were diluted five times with the buffer containing 0.01% SDS, 1% Triton X-100, 1.2 mM EDTA pH 8.0, 150 nM NaCl. 1 ug of the H3K36me2 antibody (Cell Signaling Technology, USA, Cat# 2901S) was added and incubated by rotation at 4°C overnight. Protein G Dynabeads (Life Technologies, USA) were added the next morning and incubated by rotation for an additional 4 hours. Dynabeads were next washed with 1X wash buffer followed by cold TE buffer. DNAs were reverse crosslinked, purified, followed by library preparation and deep sequencing using the Illumina HiSeq platform.

### Data sources

DNA methylome of esophageal samples were obtained from our recent work^25^, including WGBS on 21 ESCC, 3 NESQ, 5 EAC, 7 GEJ tumors and 7 NGEJ tissues. We obtained additional two NESQ samples from the ENCODE consortium to ensure statistical power. Considering the indistinguishable clinical and molecular characteristics between EAC and GEJ tumors, in the present study they were combined as the same subtype (referred to as EAC), which is a common strategy in the field^3^. TCGA Pan-cancer DNA methylome derived from HM450k methylation array was downloaded from GDC v16.0 by TCGAbiolinks package (version 2.13.6)^59^. Other DNA methylation data from individual studies, including EAC EPIC array data from the Oesophageal Cancer Clinical and Molecular Stratification (OCCAMS) consortium (EGAD00010001822)^9^, EAC and BE methylome from GSE72874^7^ and GSE81334^8^, along with ESCC tumor WGBS data (GSE149608)^6^, were analyzed for validation purposes in this study.

Other public datasets which were analyzed included: bulk ATAC-seq data of pan-cancer samples from TCGA^38^, single-cell ATAC-seq data across different adult human tissues (GSE184462)^54^, H3K27ac ChIP-seq in EAC samples (GSE132680)^39^, EAC cell lines (ESO26, FLO1, JH-EsoAd1, OACp4C, OE19, OE33, SKGT4 from GSE132680)^39^ and ESCC cell lines (KYSE140, KYSE70, TE5 from GSE106563^40^; KYSE150, KYSE180, KYSE200 from GSE131490^41^; TE7 from GSE106433^42^), HNF4A ChIP-seq in OE19 (E-MTAB-6858)^46^ and ESO26 cell lines (GSE132813)^47^, GATA4 ChIP-seq in ESO26 cell line (GSE132813)^47^ and TP63 ChIP-seq in TE5 cell line (GSE148920)^41^. H3K36me2 bigwig files of wildtype (NSD1-WT) HNSCC cell lines were downloaded from GSE149670^60^. Somatic mutation datasets were downloaded from individual studies^9,61^. We also retrieved the transcriptomic data of esophageal cancer from the TCGA consortium and GSE149609^6^.

CGI promoters are annotated as regions ranging from 250 bp upstream to 500 bp downstream of any TSSs overlapping with Takai CGIs^62^. Repetitive elements, including long interspersed nuclear elements (LINE), short interspersed nuclear elements (SINE) and long terminal repeats (LTR), were extracted from UCSC website (http://hgdownload.soe.ucsc.edu). We downloaded the annotation of common PMDs (defined as shared PMDs identified from 40 different cancer types)^19^ as well as solo-WCGW from https://zwdzwd.github.io/pmd^19^ and ENCODE blacklist regions from https://github.com/Boyle-Lab/Blacklist/tree/master/lists^63^. All of the annotations were converted to the hg38 version using the UCSC LiftOver script (https://genome.ucsc.edu/cgi-bin/hgLiftOver). The human core transcription-factor-binding sequences in the HOMOCOMO database (version 11) were used for motif annotation^64^.

### DNA methylation data analysis

For WGBS data, raw reads were mapped to the human genome (GRCh38) by Biscuit align command (version 0.1.4, https://www.githubcom/zwdzwd/biscuit) with default settings. Mapped reads were sorted by genome position, and duplicates were marked using Picard MarkDuplicates tool (version 1.136, http://broadinstitute.github.io/picard/). Biscuit pileup and vcf2bed command were then used to extract DNA methylation information. All CpG sites with a coverage >=3 informative reads and outside of the ENCODE blacklist regions were retained for downstream analyses. For EPIC and HM450K array data, methylation of each probe was extracted using the SeSAME package with noob and dyeBiasCorrTypeINorm function for background subtraction and dye bias correction^65^. According to the annotation of Infinium DNA methylation arrays^66^, recommended general masking probes were removed. HM450K methylation data were used to estimate the chromatin B compartments using minfi compartments function with “resolution=100*1000, what = “OpenSea”” options^33^.

### Development of a sequence-aware PMD calling method: Multi-model PMD SeekR (MMSeekR)

We recently performed neural network-based machine learning to establish local DNA sequence features of CpGs that were associated with global DNA methylation loss, and derived a neural network (NN) score for each CpG across the human genome^32^. In order to exclude the potential impact of high CpG density (such as CpG island), we reserved CpGs having 2 or fewer neighboring CpGs within the 151 bp window centered on the reference CpG. We investigated the correlation between NN scores and methylation in individual samples in non-overlapping 201-CpG windows across the genome. As expected, due to the greater degree of methylation loss within PMDs, there was a strong negative correlation between DNA methylation levels and NN scores within windows in PMDs, in contrast to much more modest correlations within highly methylated domains (HMD) windows (**Supplementary Fig. 1A**).

We next applied Pearson correlation coefficient (PCC) between our NN score and DNA methylation, as well as the “alpha score” used in the MethylSeekR model, to 201-CpG windows genome-wide. Compared with the NN score, the MethylSeekR alpha score is a very different measurement, returning a high score if the distribution of methylation values is closer to a unimodal beta distribution centered on 0.5 (typical of PMDs) than it is to a bimodal methylation value distribution close to 0 and 1 (typical of HMDs). Specifically, we applied a Hidden Markov Model (HMM) segmentation (as in MethylSeekR) to each model independently, and found that both the PCC and MethylSeekR alpha score showed bimodal distributions for the testing sample (**Supplementary Fig. 1B-C**). We hypothesized that since the PCC and the alpha score were very different models, combining them might improve the performance of PMD calling (**Supplementary Fig. 1D**). Thus we developed a “2-dimensional (2D)” model accordingly (**Fig. 1C**). This 2D model performed comparably well or better than either MethylSeekR or MethPipe in most cases, returning results consistently and highly overlapping with common PMDs (**Supplementary Table 2**).

While the 2D model generally performed well, we did note that it failed in a few samples with extreme methylation loss. Interestingly, these failed cases universally showed PMD methylation values very close to 0, which would be expected to violate the assumptions of both the PCC model and alpha model due to lack of variance within PMDs (**Fig. 1C right part**). We thus postulated the raw methylation values (transformed to an M-value to disperse scores close to 0 and 1) might provide additional predictive power in certain samples with extreme methylation loss, and we developed a 3D model accordingly by adding the M-value model to the 2D model. In order to decide whether the 2D or 3D model should be applied for any given sample, we first measured the methylation values of all CpGs with 2 or fewer neighboring CpGs within a 151bp window, which excludes most CpG islands, and contains a set of CpGs that are strongly associated with PMD hypomethylation^19^. If the bottom 10th percentile of these CpGs had a methylation value below 0.025, the 3D model was selected, otherwise, the 2D model was selected. This was based on the observation that the majority of samples with extreme methylation loss failed under both the MethylSeekR and MMSeekR 2D model (**Fig. 1C**).

### Application of MMSeekR to WGBS data

MMSeekR was applied to call PMDs in each WGBS sample. Before PMD calling, CpG sites with coverage of fewer than 5 informative reads were excluded. Then ENCODE blacklist regions were subtracted from the resulting PMDs. Within each esophageal cancer subtype, PMDs generated from each sample were integrated using bedtools multiinter function (version 2.27.1, https://bedtools.readthedocs.io/en/latest/). The common PMD set for each subtype contained those occurring in at least two-thirds of samples from that subtype. We further defined subtype-specific PMDs as those common PMDs from one subtype that were detected in fewer than one-third of samples in the other subtype. Meanwhile, PMDs that were in both the common EAC set and the common ESCC set were denoted as shared PMDs. Regions that were PMDs in <1/3 samples of both subtypes were denoted as shared HMDs.

### Identification and characterization of DMRs

Regions belonging to either the common ESCC or common EAC PMD sets were masked out from the DMR analysis. The Dmrseq package (version 1.10.0)^67^ was used to identify DMRs between ESCC and EAC tumors with the following parameters: cutoff =0.1, bpSpan=1000, minInSpan=30, maxPerms=500. Since the coverage information of each CpG site is required by dmrseq for statistical inference, here we included all CpG sites with >= 3 informative reads. Regions with q value < 0.05 and absolute delta methylation change > 0.2 were identified as DMRs. For hypomethylated DMRs (hypoDMRs) from each cancer subtype, we further performed one-tailed t-tests comparing the mean methylation within the DMR in nonmalignant *vs*. tumor samples, and those with FDR<0.1 were considered as tumor-specific (ts)-hypoDMRs. Both hypoDMRs and ts-hypoDMRs were annotated using HOMER annotatePeaks.pl script (version 4.9.1)^44^.

### Calculation of mean DNA methylation levels

CpG sites with a coverage of at least 5 informative reads were used for this calculation. Average methylation levels of CpG sites across the genome (global level), within CGI promoters, commonPMDs, SINE, LINE and LTR in each sample were calculated independently. Besides, we obtained the mean methylation of CpG sites in non-PMD regions. For genome/domain-wide visualization, the average methylation of 10-kb consecutive non-overlapping tiles was shown. To calculate the mean methylation levels within shared PMDs/HMDs, EAC-specific PMDs and ESCC-specific PMDs, solo-WCGW CpG sites/probes were used.

### Principal component analysis of WGBS data

PMDs were identified by either MethPipe, MethylSeekR or MMseekR (**Fig. 1D**). The whole genome was split into 30-kb consecutive but non-overlapping tiles. For each tile, the ratio overlapping with any PMD was calculated for each caller. The top 5,000 most variable 30-kb tiles from each PMD caller were used in Principal component analysis (PCA). In **Supplementary Fig. 3A** and **3B**, CpG sites with at least 7 reads across all esophageal samples were used. Then the top 8,000 most variable CpG sites were selected for PCA using the R prcomp function. PCA was performed before and after masking the combined common PMDs from EAC and ESCC, and generated the point plots by ggplot2 package (version 3.1.0).

### RNA-seq data analysis

According to the raw read counts obtained from the TCGA, we identified significant upregulated genes by DESeq2 package (version 1.22.2) with adjusted p-value < 0.05, fold change > 1.5 and mean FPKM >1 in the corresponding sample groups^68^. For expression datasets of nonmalignant squamous and ESCC tissues, raw reads were aligned to GRCh38 using HISAT2 (version 2.0.4)^69^ and quantified by htseq-count program (version 0.11.2) at default setting. Significant upregulated genes were identified using the same method as for the TCGA datasets.

### ChIP-seq data analysis

Raw reads were mapped to GRCh38 (ENSEMBL release 84) using BWA mem program (version 0.7.15) with the default options^70^. Then the mapped reads were sorted using SAMtools program (version 1.3.1)^71^, followed by removing PCR duplicates and blacklist regions by Picard MarkDuplicates tool and bedtools (version 2.27.1). MACS2 (Model-Based Analysis of ChIP-Seq, version 2.1.2) were applied to call peaks with the default setting for transcription factors, ‘‘-q 0.01-extsize = 146 −nomodel’’ options for H3K27ac and ‘‘--broad -p 0.01 --extsize=146 --nomodel’’ for H3K36me2^72^. Bigwig files were generated by deepTools bamCompare function (version 3.1.3) with “--operation subtract --normalizeUsing CPM --extendReads 146 --binSize 20” parameters^73^. Average signals of shared PMDs/HMDs, EAC-only PMDs and ESCC-only PMDs in each H3K27ac or H3K36me2 ChIP-seq sample were extracted from bigwig files using deepTools computeMatrix function with ‘‘scale-regions’’ option.

### ATAC-seq data analysis

For bulk pan-cancer ATAC-seq data obtained from the TCGA project, the average accessibility of regions/domains was extracted from the available bigwig files using deepTools computeMatrix function^38^. To avoid the influence of scaling factors across different samples and batches, the mean accessibility across the whole genome in each sample was calculated and used for normalization. For single cell ATAC-seq data, based on the clustering and annotation results from the publication^54^, only epithelial cell types were used for further analysis. Similarly, the average accessibility of regions/domains was derived for each cell in each sample and normalized by the mean signal across the whole genome.

### DMR motif enrichment analysis

For each hypoDMR or ts-hypoDMR, we randomly sampled 10 regions with the same size and number of CpGs to define the background set. Then motif searching of both DMRs and background regions was performed using HOMER annotatePeaks.pl with ‘‘-noann -m HOCOMOCOv11_core_HUMAN_mono_homer_format_0.0001.motif’’ parameters^44^. The ELMER method was next applied to identify potential transcription-factor-binding sequences and the top 15 transcription factors with q-value < 0.05 and FPKM > 5 in the corresponding cancer subtype were reserved for further analysis^74^.

### Pathway enrichment analysis

We performed the pathway (Biological Process) enrichment analysis by Cistrome-GO^75^ using candidate regions with methylation changes and differential expression analysis results. For hypoDMR analysis, subtype-specific DMRs and upregulated genes in the corresponding tumors were used as input data. For subtype-specific PMDs, the input data contained PMD regions and downregulated genes in the corresponding tumors. The top 15 enriched pathways with q value < 0.05 were shown.

## Code Availability

Source code for MMSeekR is available at https://github.com/yuanzi2/MMSeekR. Source code for WGBS data analysis and figure reproduction is in https://github.com/yuanzi2/ESCA_WGBS_analysis.

## Data Availability

WGBS data and ChIP-seq data for H3K36me2 in EAC and ESCC cell lines were available at GSE210220.

## Acknowledgement

We thank the OCCAMS Study for sharing DNA methylation and somatic mutation data of EAC samples. D-C.L. was supported by NIH/NCI under awards P30CA014089 and R37CA237022. Y.Y.Z. was partially supported by the Fundamental Research Funds For the Central Universities, Sun Yat-sen University (22qntd3701). This work is also partially funded by the institutional funds from the Herman Ostrow School of Dentistry of USC’s Center for Craniofacial Molecular Biology to B.Z and D-C.L, and a Project Grant (845755) from the Israel Cancer Research Fund Project Grant to B.P.B.

## Author contribution

D.-C.L. and B.P.B. conceived and devised the study. D.-C.L., B.P.B., Y.Y.Z., and B.Z. designed experiments and analyses. Y.Y.Z and B.P.B. performed bioinformatics and statistical analysis. B.Z performed the experiments. Y.Y.Z., B.P.B., and D.-C.L. analyzed the data. B.P.B., D.-C.L. supervised the research. A.S.H, U.K.S, L.Y.X, E.M.L and H.P.K. contributed the data and materials. Y.Y.Z., and D.-C.L. wrote the manuscript with input from B.P.B. The last two authors (D.-C.L. and B.P.B.) are co-senior authors who jointly supervised the work, and they have the right to list their names last in their CV.

## Supplementary information

**Supplementary Figures.docx**

**Supplementary Table 1.** WGBS data sets used in the current study.

**Supplementary Table 2.** The percent of PMDs identified by three different callers overlapping with common PMDs or HMDs in each tumor sample from the Blueprint consortium or esophageal tissue.

## Reference

1. Sung, H. et al. Global Cancer Statistics 2020: GLOBOCAN Estimates of Incidence and Mortality Worldwide for 36 Cancers in 185 Countries. CA Cancer J. Clin. 71, 209–249 (2021).

2. Siegel, R. L., Miller, K. D., Fuchs, H. E. & Jemal, A. Cancer Statistics, 2021. CA Cancer J. Clin. 71, 7–33 (2021).

3. Cancer Genome Atlas Research Network et al. Integrated genomic characterization of oesophageal carcinoma. Nature 541, 169–175 (2017).

4. Talukdar, F. R. et al. Genome-Wide DNA Methylation Profiling of Esophageal Squamous Cell Carcinoma from Global High-Incidence Regions Identifies Crucial Genes and Potential Cancer Markers. Cancer Res. 81, 2612–2624 (2021).

5. Teng, H. et al. Inter- and intratumor DNA methylation heterogeneity associated with lymph node metastasis and prognosis of esophageal squamous cell carcinoma. Theranostics 10, 3035–3048 (2020).

6. Cao, W. et al. Multi-faceted epigenetic dysregulation of gene expression promotes esophageal squamous cell carcinoma. Nat. Commun. 11, 3675 (2020).

7. Krause, L. et al. Identification of the CIMP-like subtype and aberrant methylation of members of the chromosomal segregation and spindle assembly pathways in esophageal adenocarcinoma. Carcinogenesis 37, 356–365 (2016).

8. Yu, M. et al. Subtypes of Barrett’s oesophagus and oesophageal adenocarcinoma based on genome-wide methylation analysis. Gut 68, 389–399 (2019).

9. Jammula, S. et al. Identification of Subtypes of Barrett’s Esophagus and Esophageal Adenocarcinoma Based on DNA Methylation Profiles and Integration of Transcriptome and Genome Data. Gastroenterology 158, 1682–1697.e1 (2020).

10. Angeloni, A. & Bogdanovic, O. Enhancer DNA methylation: implications for gene regulation. Essays Biochem. 63, 707–715 (2019).

11. Lister, R. et al. Human DNA methylomes at base resolution show widespread epigenomic differences. Nature 462, 315–322 (2009).

12. Slotkin, R. K., Keith Slotkin, R. & Martienssen, R. Transposable elements and the epigenetic regulation of the genome. Nature Reviews Genetics vol. 8 272–285 (2007).

13. Baylin, S. B. & Jones, P. A. Epigenetic Determinants of Cancer. Cold Spring Harb. Perspect. Biol. 8, (2016).

14. Luo, C., Hajkova, P. & Ecker, J. R. Dynamic DNA methylation: In the right place at the right time. Science 361, 1336–1340 (2018).

15. Karlow, J. A., Miao, B., Xing, X., Wang, T. & Zhang, B. Common DNA methylation dynamics in endometriod adenocarcinoma and glioblastoma suggest universal epigenomic alterations in tumorigenesis. Commun Biol 4, 607 (2021).

16. Hansen, K. D. et al. Increased methylation variation in epigenetic domains across cancer types. Nat. Genet. 43, 768–775 (2011).

17. Berman, B. P. et al. Regions of focal DNA hypermethylation and long-range hypomethylation in colorectal cancer coincide with nuclear lamina-associated domains. Nat. Genet. 44, 40–46 (2011).

18. Hon, G. C. et al. Global DNA hypomethylation coupled to repressive chromatin domain formation and gene silencing in breast cancer. Genome Res. 22, 246–258 (2012).

19. Zhou, W. et al. DNA methylation loss in late-replicating domains is linked to mitotic cell division. Nat. Genet. 50, 591–602 (2018).

20. Duran-Ferrer, M. et al. The proliferative history shapes the DNA methylome of B-cell tumors and predicts clinical outcome. Nat Cancer 1, 1066–1081 (2020).

21. Hur, K. et al. Hypomethylation of long interspersed nuclear element-1 (LINE-1) leads to activation of proto-oncogenes in human colorectal cancer metastasis. Gut 63, 635–646 (2014).

22. Hovestadt, V. et al. Decoding the regulatory landscape of medulloblastoma using DNA methylation sequencing. Nature 510, 537–541 (2014).

23. Brinkman, A. B. et al. Partially methylated domains are hypervariable in breast cancer and fuel widespread CpG island hypermethylation. Nat. Commun. 10, 1749 (2019).

24. Salhab, A. et al. A comprehensive analysis of 195 DNA methylomes reveals shared and cell-specific features of partially methylated domains. Genome Biol. 19, 150 (2018).

25. Pan, F. et al. Characterization of epigenetic alterations in esophageal cancer by whole-genome bisulfite sequencing. bioRxiv 2021.12.05.471340 (2021) doi:10.1101/2021.12.05.471340.

26. Liu, Y. et al. Comparative Molecular Analysis of Gastrointestinal Adenocarcinomas. Cancer Cell 33, 721–735.e8 (2018).

27. Tao, Y. et al. Aging-like Spontaneous Epigenetic Silencing Facilitates Wnt Activation, Stemness, and Braf-Induced Tumorigenesis. Cancer Cell 35, 315–328.e6 (2019).

28. Vaz, M. et al. Chronic Cigarette Smoke-Induced Epigenomic Changes Precede Sensitization of Bronchial Epithelial Cells to Single-Step Transformation by KRAS Mutations. Cancer Cell 32, 360–376.e6 (2017).

29. Ehrlich, M. & Lacey, M. DNA hypomethylation and hemimethylation in cancer. Adv. Exp. Med. Biol. 754, 31–56 (2013).

30. Decato, B. E. et al. Characterization of universal features of partially methylated domains across tissues and species. Epigenetics Chromatin 13, 39 (2020).

31. Burger, L., Gaidatzis, D., Schübeler, D. & Stadler, M. B. Identification of active regulatory regions from DNA methylation data. Nucleic Acids Res. 41, e155 (2013).

32. Bar, D. et al. A local sequence signature defines a subset of heterochromatin-associated CpGs with minimal loss of methylation in healthy tissues but extensive loss in cancer. bioRxiv 2022.08.16.504069 (2022) doi:10.1101/2022.08.16.504069.

33. Fortin, J.-P. & Hansen, K. D. Reconstructing A/B compartments as revealed by Hi-C using long-range correlations in epigenetic data. Genome Biol. 16, 180 (2015).

34. Schuster-Böckler, B. & Lehner, B. Chromatin organization is a major influence on regional mutation rates in human cancer cells. Nature 488, 504–507 (2012).

35. Lawrence, M. S. et al. Mutational heterogeneity in cancer and the search for new cancer-associated genes. Nature 499, 214–218 (2013).

36. Weinberg, D. N. et al. The histone mark H3K36me2 recruits DNMT3A and shapes the intergenic DNA methylation landscape. Nature 573, 281–286 (2019).

37. Neri, F. et al. Intragenic DNA methylation prevents spurious transcription initiation. Nature 543, 72–77 (2017).

38. Corces, M. R. et al. The chromatin accessibility landscape of primary human cancers. Science 362, (2018).

39. Chen, L. et al. Master transcription factors form interconnected circuitry and orchestrate transcriptional networks in oesophageal adenocarcinoma. Gut 69, 630–640 (2020).

40. Jiang, Y. et al. Co-activation of super-enhancer-driven CCAT1 by TP63 and SOX2 promotes squamous cancer progression. Nat. Commun. 9, 3619 (2018).

41. Jiang, Y.-Y. et al. TP63, SOX2, and KLF5 Establish a Core Regulatory Circuitry That Controls Epigenetic and Transcription Patterns in Esophageal Squamous Cell Carcinoma Cell Lines. Gastroenterology 159, 1311–1327.e19 (2020).

42. Xie, J.-J. et al. Super-Enhancer-Driven Long Non-Coding RNA LINC01503, Regulated by TP63, Is Over-Expressed and Oncogenic in Squamous Cell Carcinoma. Gastroenterology 154, 2137–2151.e1 (2018).

43. Espinet, E. et al. Aggressive PDACs Show Hypomethylation of Repetitive Elements and the Execution of an Intrinsic IFN Program Linked to a Ductal Cell of Origin. Cancer Discov. 11, 638–659 (2021).

44. Heinz, S. et al. Simple combinations of lineage-determining transcription factors prime cis-regulatory elements required for macrophage and B cell identities. Mol. Cell 38, 576–589 (2010).

45. Aran, D., Sabato, S. & Hellman, A. DNA methylation of distal regulatory sites characterizes dysregulation of cancer genes. Genome Biol. 14, R21 (2013).

46. Rogerson, C. et al. Identification of a primitive intestinal transcription factor network shared between esophageal adenocarcinoma and its precancerous precursor state. Genome Res. 29, 723–736 (2019).

47. Pan, J. et al. Lineage-Specific Epigenomic and Genomic Activation of Oncogene HNF4A Promotes Gastrointestinal Adenocarcinomas. Cancer Res. 80, 2722–2736 (2020).

48. Lopez-Pajares, V. et al. A LncRNA-MAF:MAFB transcription factor network regulates epidermal differentiation. Dev. Cell 32, 693–706 (2015).

49. Reddy, J. et al. Predicting master transcription factors from pan-cancer expression data. Sci Adv 7, eabf6123 (2021).

50. Sanda, T. et al. Core transcriptional regulatory circuit controlled by the TAL1 complex in human T cell acute lymphoblastic leukemia. Cancer Cell 22, 209–221 (2012).

51. Walker, E. M., Thompson, C. A. & Battle, M. A. GATA4 and GATA6 regulate intestinal epithelial cytodifferentiation during development. Dev. Biol. 392, 283–294 (2014).

52. Ye, D. Z. & Kaestner, K. H. Foxa1 and Foxa2 control the differentiation of goblet and enteroendocrine L- and D-cells in mice. Gastroenterology 137, 2052–2062 (2009).

53. Britton, E. et al. Open chromatin profiling identifies AP1 as a transcriptional regulator in oesophageal adenocarcinoma. PLoS Genet. 13, e1006879 (2017).

54. Zhang, K. et al. A single-cell atlas of chromatin accessibility in the human genome. Cell 184, 5985–6001.e19 (2021).

55. Hoadley, K. A. et al. Cell-of-Origin Patterns Dominate the Molecular Classification of 10,000 Tumors from 33 Types of Cancer. Cell 173, 291–304.e6 (2018).

56. Nothjunge, S. et al. DNA methylation signatures follow preformed chromatin compartments in cardiac myocytes. Nat. Commun. 8, 1667 (2017).

57. Du, Q. et al. DNA methylation is required to maintain both DNA replication timing precision and 3D genome organization integrity. Cell Rep. 36, 109722 (2021).

58. Johnstone, S. E. et al. Large-Scale Topological Changes Restrain Malignant Progression in Colorectal Cancer. Cell 182, 1474–1489.e23 (2020).

59. Mounir, M. et al. New functionalities in the TCGAbiolinks package for the study and integration of cancer data from GDC and GTEx. PLoS Comput. Biol. 15, e1006701 (2019).

60. Farhangdoost, N. et al. Chromatin dysregulation associated with NSD1 mutation in head and neck squamous cell carcinoma. Cell Rep. 34, 108769 (2021).

61. Cui, Y. et al. Whole-genome sequencing of 508 patients identifies key molecular features associated with poor prognosis in esophageal squamous cell carcinoma. Cell Res. 30, 902–913 (2020).

62. Takai, D. & Jones, P. A. Comprehensive analysis of CpG islands in human chromosomes 21 and 22. Proc. Natl. Acad. Sci. U. S. A. 99, 3740–3745 (2002).

63. Amemiya, H. M., Kundaje, A. & Boyle, A. P. The ENCODE Blacklist: Identification of Problematic Regions of the Genome. Sci. Rep. 9, 9354 (2019).

64. Kulakovskiy, I. V. et al. HOCOMOCO: towards a complete collection of transcription factor binding models for human and mouse via large-scale ChIP-Seq analysis. Nucleic Acids Res. 46, D252–D259 (2018).

65. Zhou, W., Triche, T. J., Jr, Laird, P. W. & Shen, H. SeSAMe: reducing artifactual detection of DNA methylation by Infinium BeadChips in genomic deletions. Nucleic Acids Res. 46, e123 (2018).

66. Zhou, W., Laird, P. W. & Shen, H. Comprehensive characterization, annotation and innovative use of Infinium DNA methylation BeadChip probes. Nucleic Acids Res. 45, e22 (2017).

67. Korthauer, K., Chakraborty, S., Benjamini, Y. & Irizarry, R. A. Detection and accurate false discovery rate control of differentially methylated regions from whole genome bisulfite sequencing. Biostatistics 20, 367–383 (2019).

68. Love, M. I., Huber, W. & Anders, S. Moderated estimation of fold change and dispersion for RNA-seq data with DESeq2. Genome Biol. 15, 550 (2014).

69. Kim, D., Langmead, B. & Salzberg, S. L. HISAT: a fast spliced aligner with low memory requirements. Nat. Methods 12, 357–360 (2015).

70. Li, H. Aligning sequence reads, clone sequences and assembly contigs with BWA-MEM. (2013).

71. Li, H. et al. The Sequence Alignment/Map format and SAMtools. Bioinformatics 25, 2078–2079 (2009).

72. Zhang, Y. et al. Model-based analysis of ChIP-Seq (MACS). Genome Biol. 9, R137 (2008).

73. Ramírez, F. et al. deepTools2: a next generation web server for deep-sequencing data analysis. Nucleic Acids Res. 44, W160–5 (2016).

74. Silva, T. C. et al. ELMER v.2: an R/Bioconductor package to reconstruct gene regulatory networks from DNA methylation and transcriptome profiles. Bioinformatics 35, 1974–1977 (2019).

75. Li, S. et al. Cistrome-GO: a web server for functional enrichment analysis of transcription factor ChIP-seq peaks. Nucleic Acids Res. 47, W206–W211 (2019).

76. Irizarry, R. A. et al. Genome-wide methylation analysis of human colon cancer reveals similar hypo- and hypermethylation at conserved tissue-specific CpG island shores. Nat. Genet. 41, 178 (2009).

77. Silva, T. C. et al. ELMER v.2: An R/Bioconductor package to reconstruct gene regulatory networks from DNA methylation and transcriptome profiles. doi:10.1101/148726.

