## Supplementary Figures for "Comprehensive analyses of partially methylated domains and differentially methylated regions in esophageal cancer reveal both cell-type- and cancer-specific epigenetic regulation"

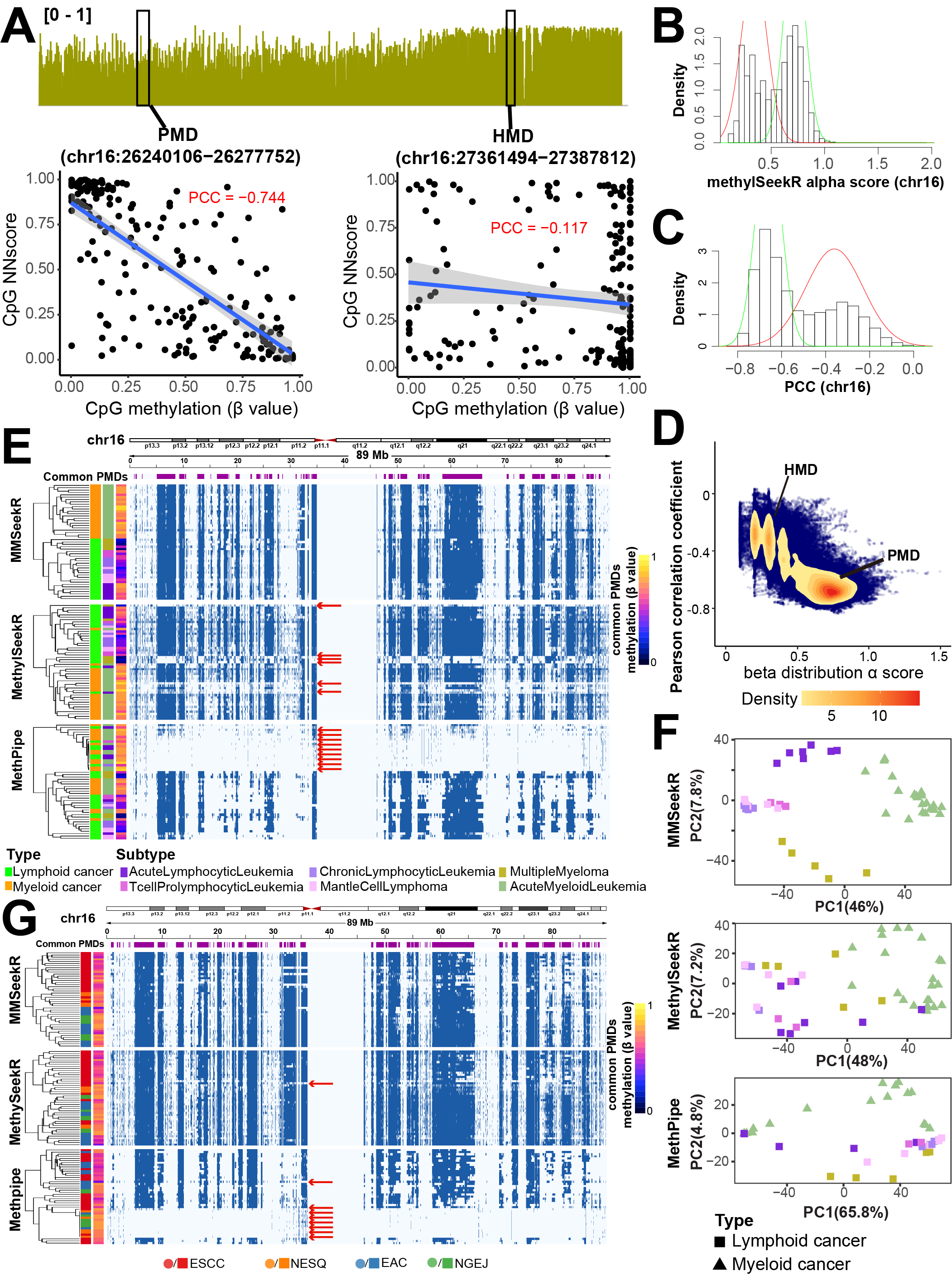

**Supplementary Figure 1. The** **development of MMSeekR, a sequence-aware multi-model PMD caller.** **(A)** Pearson correlation coefficients between NN scores and DNA methylation levels in individual CpGs in representative PMD or HMD windows. **(B)** MethylSeekR α scores and (**C**) Pearson correlation coefficient across all 201-CpG windows of chr16 show bimodal distributions. **(D)** A density plot of the distributions of Pearson correlation coefficient and MethylSeekR α scores. The methylome data in **(A-D)** are from a Blueprint tumor sample “S01FJZA1_MantleCellLymphoma”. **(E)** PMD regions (blue) across chr16 identified by the three PMD callers; the 47 Blueprint samples were clustered by the PMD distribution across the genome. Red arrows point to failures in MethylSeekR and Methpipe methods. **(F)** PCA analysis using the top 5,000 most variable 30-kb tiles from the three PMD callers. Data used in this figure are from Blueprint consortium. **(G)** Similar to (**D)**, showing PMD regions (blue) across chr16 identified by the three PMD callers; esophageal samples were clustered by PMD distribution across the genome.

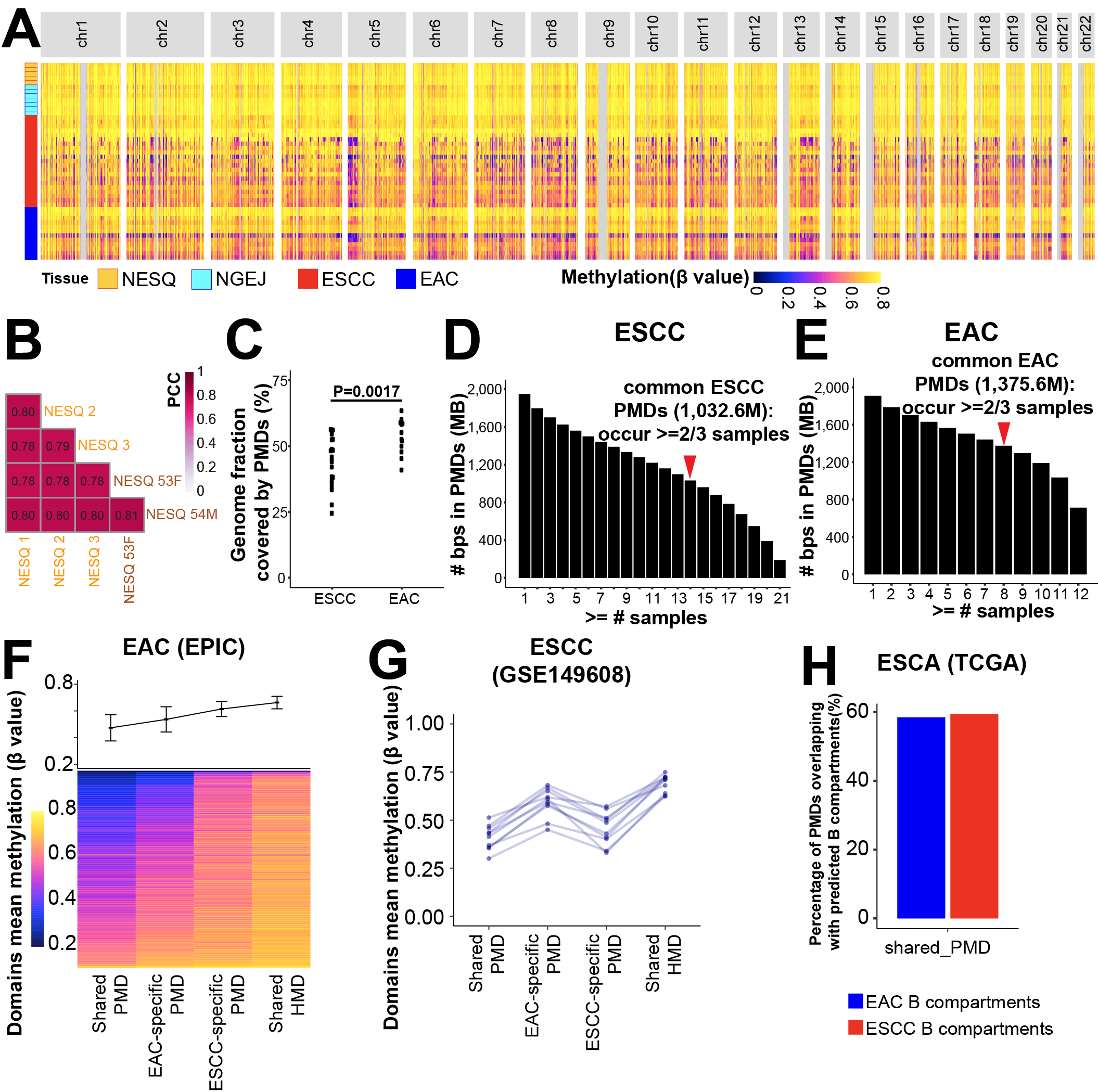

**Supplementary Figure 2. Analyses of subtype-specific PMDs. (A)** Genome-wide maps of DNA methylation profiles from 45 esophageal WGBS samples. Average methylation values were shown in consecutive and non-overlapping 10-kb tiles. CGI regions were masked using the annotation from Irizarry et al. **(B)** NESQ tissues show high inter-sample correlation in DNA methylation levels despite from two different datasets. NESQ1, NESQ2 and NESQ3 are internal samples while the other two are from ENCODE projects. **(C)** Genome fractions covered by PMDs in each tumor sample. The P value was determined by a two-tail t test. **(D-E)** Genomic regions covered by PMDs which are common in ESCC **(D)** or EAC samples **(E)**. **(F)** Heatmaps and line plots displaying the methylation levels for different PMD categories in EAC. Each row in the heatmap shows methylation (beta value) per sample; the trend line displays the average methylation and standard deviation. **(G)** Line plots showing the average methylation levels for different PMD categories in ESCC. Each line represents one sample. **(H)** Bar plots showing the percentage of shared PMDs overlapping with chromatin B compartments, which were defined by TCGA methylation datasets using minfi package.

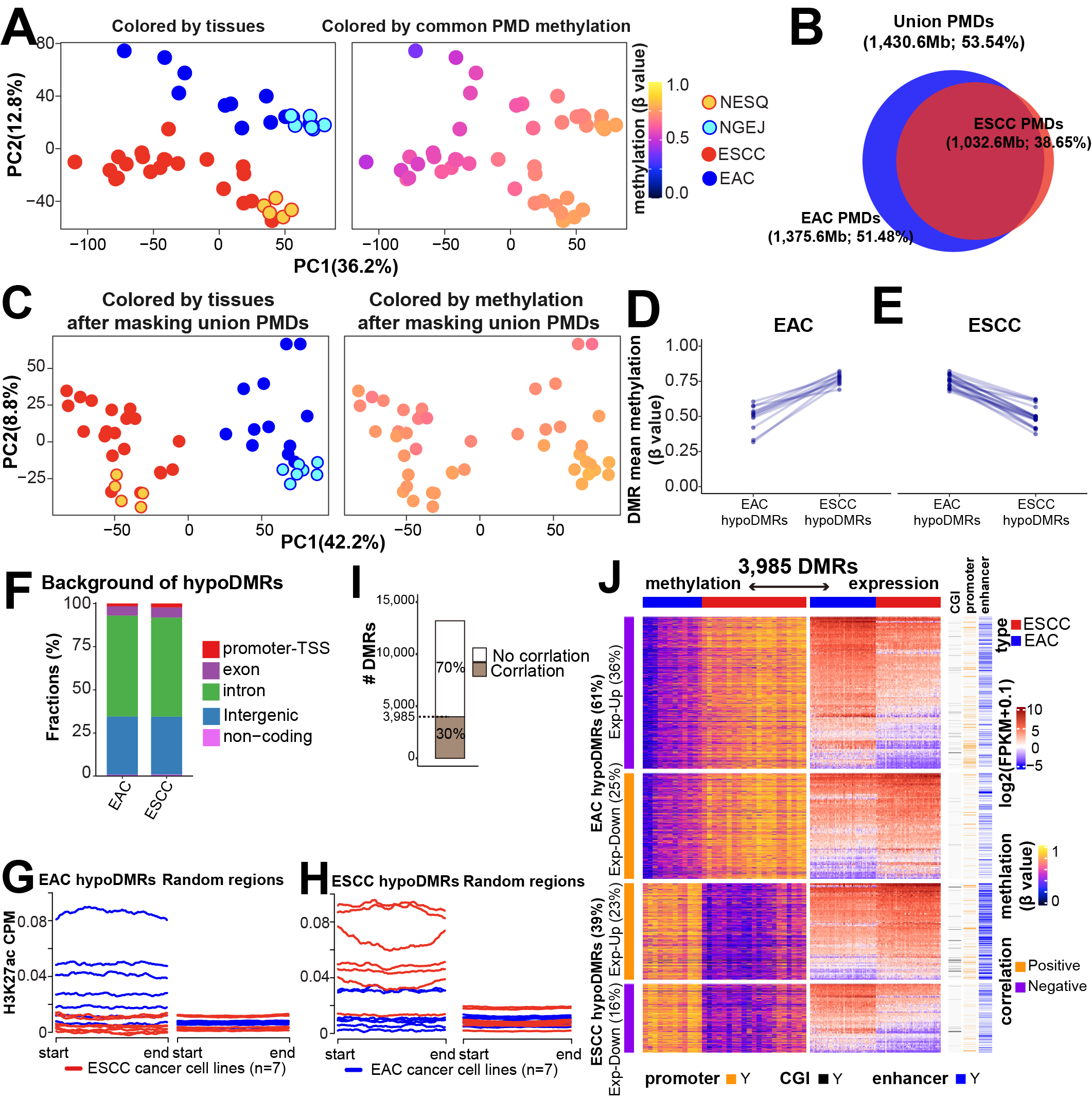

**Supplementary Figure 3. DMR analyses upon masking of union PMDs. (A)** PCA analysis using the top 8,000 most variable CpGs with a coverage higher than 7 in each sample. The samples are colored by tissue types (left) and methylation levels of common PMDs (right). **(B)** A Venn diagram showing the union PMD set combining EAC and ESCC PMDs, which were identified according to **Supplementary Figure 2B-C**. **(C)** PCA analysis upon masking the union PMD set, again using the top 8,000 most variable CpG sites with a coverage higher than 7. **(D-E)** Line plots showing the average methylation levels for different hypoDMR categories in EAC **(D)** and ESCC **(E)**. Each line represents one sample. **(F)** Stacked bar plots showing fractions of hypoDMRs that overlap with different genomic features. Random genomic regions contained 10-times randomly selected regions with the same CpG density. **(G-H)** Aggregation line plots showing H3K27ac ChIP-seq signals in either EAC **(G)** or ESCC **(H)** hypoDMRs from esophageal cancer cell lines. **(I)** Number of DMRs correlated with gene expression levels. **(J)** Heatmaps showing DMRs significantly correlated with gene expression levels. The left heatmap shows the methylation level (beta value) per DMR per sample; the right heatmap displays the mRNA level of the closest gene to the DMR per sample (z-scored). DMRs overlapping with different genomic features were shown. Methylation values were extracted from the WGBS dataset. Expression data were obtained from the TCGA project.

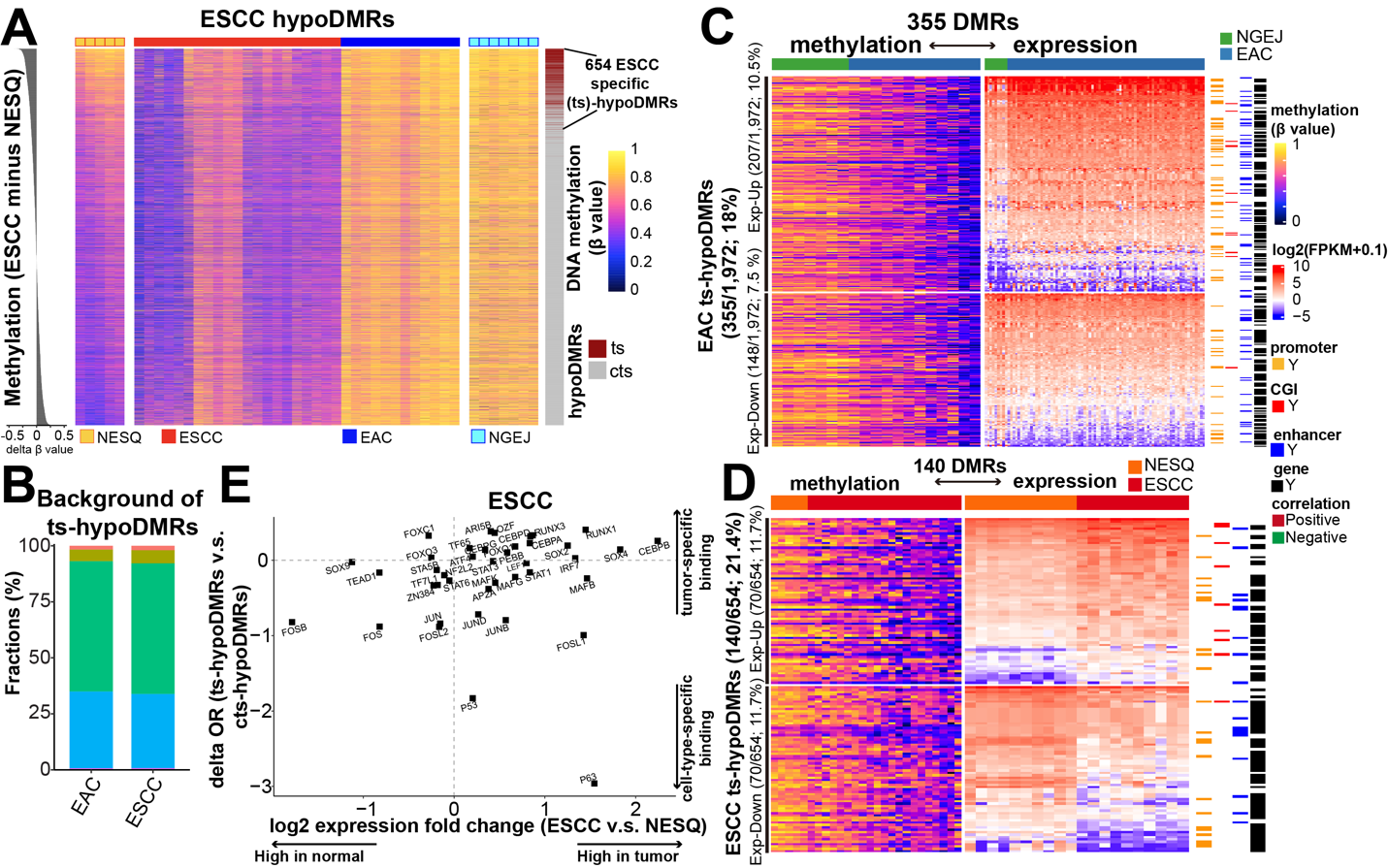

**Supplementary Figure 4. Characterization of tumor-specific hypoDMRs. (A)** Similar to **Fig. 6A**, heatmaps showing methylation levels for each ESCC hypoDMR. 654 ESCC ts-hypoDMRs were identified using one-tailed t test between ESCC and NESQ samples (right) with the FDR cutoff < 0.05. **(B)** Stacked bar plots showing fractions of ts-hypoDMRs associated with random regions that overlap with different genomic features. **(C-D)** Heatmaps showing EAC **(C)** or ESCC **(D)** ts-hypoDMRs correlated with gene expression levels. The left heatmap shows the methylation level (beta value) per DMR per sample; the right heatmap shows the mRNA expression level of the closest gene to the DMR per sample (z-scored). DMRs overlapping with different genomic features are shown. Methylation values were extracted from the WGBS dataset. The expression of EAC and paired nonmalignant tissues were obtained from the TCGA project, while the ESCC expression data were from GSE149609. **(E)** Scatter plots showing the transcription-factor-binding sites that were enriched in ESCC ts-hypoDMRs compared with cts-hypoDMRs. The X axis represents the expression fold change between ESCC and paired NESQ samples. The Y axis shows the delta enrichment score of transcription-factor-binding sites between ts- *vs.* cts-hypoDMRs. Expression data were from the GSE149609 and motif enrichment analyses were performed by ELMER.

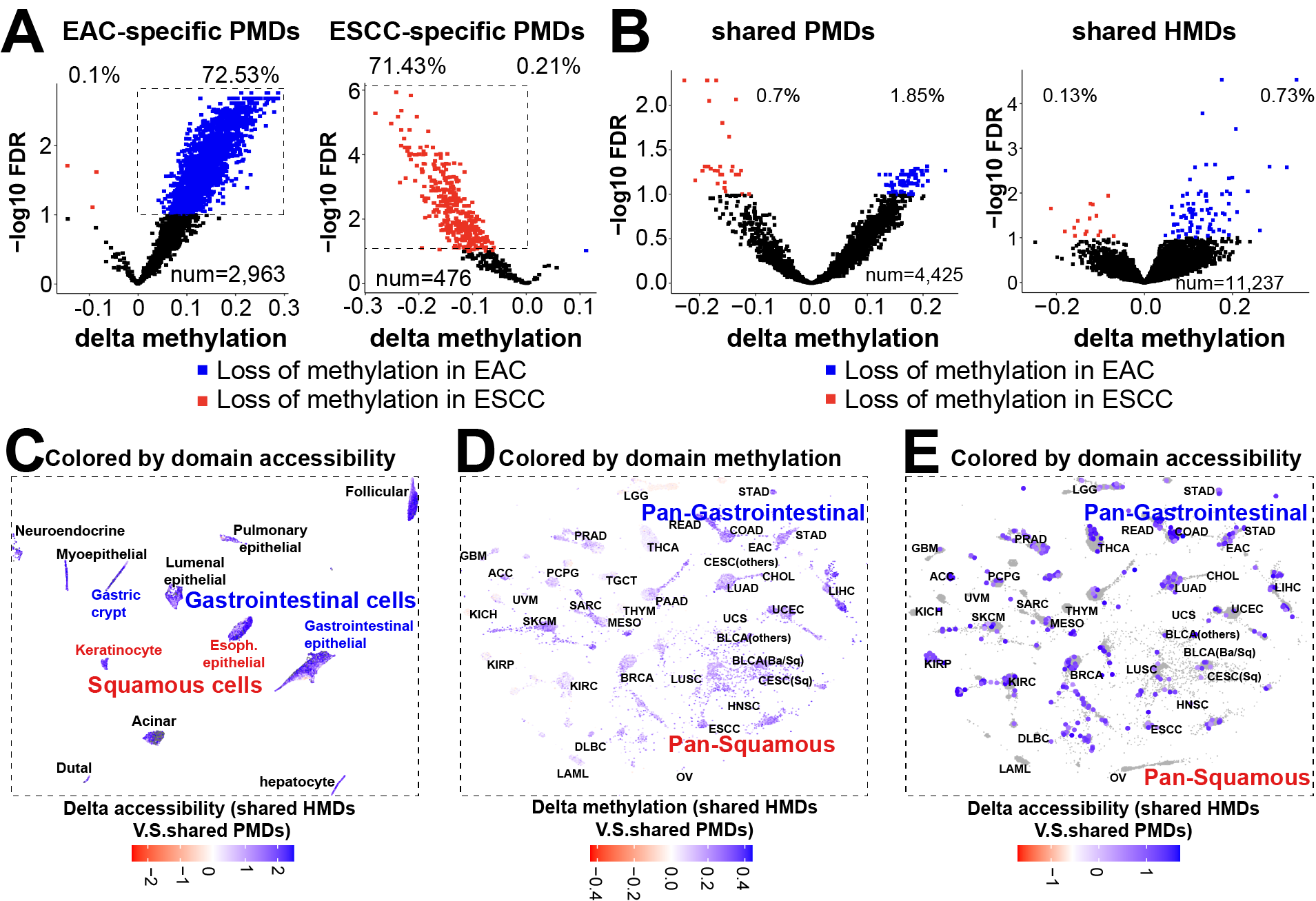

**Supplementary Figure 5. PMDs and hypoDMRs harbor prominent cell-type specificity. (A-B)** Volcano plots showing average methylation levels for different PMD categories comparing EAC *vs.* ESCC samples. The significance was determined by two-tailed t test with the FDR cutoff < 0.1 **(C)** UMAP plots showing normal single-cell clusters colored by the delta ATAC-seq accessibility between shared HMDs and PMDs. The total cell number is 145,594. **(D-E)** Pan-cancer clusters are colored by delta methylation **(D)** or ATAC-seq accessibility **(E)** between shared HMDs and PMDs. **(D)** and **(E)** contain 8,915 and 365 tumor samples, respectively.
